# Signatures of selective sweeps in urban and rural white clover populations

**DOI:** 10.1101/2024.10.30.621108

**Authors:** James S. Santangelo, Marc T. J. Johnson, Rob W. Ness

## Abstract

Urbanization is increasingly recognized as a powerful force of evolutionary change. However, anthropogenic sources of selection can often be similarly strong and mul­ tifarious in rural habitats, and whether selection differs in either strength or its targets between habitats is rarely considered. Despite numerous examples of phe­ notypic differentiation between urban and rural populations, we still lack an un­ derstanding of the genes enabling adaptation to these contrasting habitats. In this study, we conducted whole genome sequencing of 120 urban, suburban, and rural white clover plants from Toronto, Canada and used these data to identify urban and rural signatures of positive selection. We found evidence for selection in genomic regions involved in abiotic stress tolerance and growth/ development in both urban and rural populations, and clinal change in allele frequencies at SNPs within these regions. Patterns of allele frequency and haplotype differentiation suggest that most sweeps are incomplete and our strongest signals of selective sweeps overlap known large-effect structural variants. These results highlight how both urban and rural habitats are driving ongoing selection in Toronto white clover populations, and mo­ tivate future work disentangling the genetic architecture of ecologically important phenotypes underlying adaptation to contemporary anthropogenic habitats.

## Introduction

Uncovering the genetic basis and architecture of adaptation to rapid environmental change remains a central goal in evolutionary biology. Urban environments are emerging as “living laboratories” for investigations of contemporary evolution, owing in part to their global replication and their pervasive environmental impacts. These pronounced and repeated changes have the potential to drive rapid ecological and evolutionary change in natural populations (Diamond and Martin 2021; Johnson and Munshi-South 2017; Santangelo et al. 2018; Szulkin et al. 2020; Verrelli et al. 2022). While altered patterns of gene flow and genetic drift are some of the most frequently cited examples of urban evolutionary change (Carlen and Munshi-South 2021; Fusco et al. 2024; Harris et al. 2016; Lourern;o et al. 2017; Miles et al. 2019; Mueller et al. 2018; Munshi-South 2012; Schmidt et al. 2020), there is growing evidence of adaptive phenotypic responses to urbanization. For example, elevated temperatures within cities (i.e., the urban heat island effect) have led to increased thermal tolerance and changes in life-history strategies (Brans et al. 2018; Brans and De Meester 2018; Brans et al. 2017; Diamond et al. 2017, 2018; Martin et al. 2021), while urban structures have driven changes in animal morphology, locomotion, and behaviour (Evans et al. 2012; Winchell et al. 2018; Winchell et al. 2017, 2016). In some cases, the genetic basis underlying phenotypic responses to urbanization are known-for example, the two-locus epistasis underlying urban-rural cyanogenesis dines in white clover (Santangelo et al. 2022b), or regulatory changes to the Aryl hydrocarbon receptor **(AHR)** signaling pathway facilitating resistance of Gulf and Atlantic Killifish populations to toxic urban pollutants (Oziolor et al. 2019; Reid et al. 2016). Despite these examples, most studies have focused on candidate genes or reduced-representation sequencing (e.g., GBS) and thus we still lack a general understanding of the genetic architecture and genome-wide responses underlying adaptation to urban environments, and the strength and timescale of selection over which adaptation has occurred.

During the initial phases of adaptation to a new phenotypic optimum, rapid adaptation is predicted to occur primarily via changes in loci of large effect (Hay­ ward and Sella 2022). This is likely to be the case is urban environments given the substantial ecological changes they often impose. In addition, the relatively young age of cities suggests that most adaptation is likely to occur via selection on standing genetic variation, as new mutations are unlikely to have had time to accumulate. Genome-wide scans of natural selection-for example, using distortions in the site­ or haplotype-frequency spectrum, or identifying regions of extended haplotype ho­ mozygosity generated by positive selection-and gene-environment association anal­ yses (GEAs), can help generate hypotheses about putatively adaptive phenotypes, the selective agents underlying adaptive differentiation, and the number and effect sizes of alleles underlying local adaptation (Hoban et al. 2016; Lasky et al. 2023; Weigand and Leese 2018). In an urban context, recent work has identified signa­ tures of selection among urban populations in genes related to metabolism and diet (Harpak et al. 2021; Harris and Munshi-South 2017; Ravinet et al. 2018), neural function and cognition (Mueller et al. 2020; Salmon et al. 2021), thermal physiology and environmental stress response (Campbell-Staton et al. 2020; Theodorou et al. 2018), and tolerance to pollution (Kamdem et al. 2017; Oziolor et al. 2019). While the number of studies investigating molecular adaptation to urban environments is rapidly increasing, none have compared these signatures to those of nearby rural populations.

Studies of urban adaptation frequently compare the phenotypic or genomic re­ sponses of urban populations to those of nonurban, often rural, populations. Such studies are therefore primed for investigating urban-rural differences in the genetic basis and architecture of adaptation, though rural-specific signatures of selection are often ignored. Croplands and pastures currently account for approximately 40% of earth’s ice-free land (Ellis and Ramankutty 2008; Foley et al. 2005), and anthro­ pogenic sources of selection in these habitats can be strong and multifarious, often rivaling those observed in urban environments. Indeed, agricultural landscapes have long been recognized as potential hotbeds of ecological changes that can drive the evolution of contemporary populations (Baker 1974; Lau and Funk 2023; Neve et al. 2009; Snaydon and Davies 1972; Turcotte et al. 2017). For example, widespread herbicide use has driven strong and rapid increases in resistance alleles among agri­ culturally problematic weeds over just the past few decades (Kreiner et al. 2022a,b). While the agents and targets of selection might differ between urban and rural habi­ tats, ignoring signatures of adaptation in rural habitats risks missing an important source of anthropogenic selection and giving the misleading impression that the bulk of contemporary adaptation to anthropogenic disturbances is happening in cities, rather than the agricultural landscapes that support them.

In this paper, we perform genome-wide scans for signatures of selective sweeps to identify loci putatively involved in adaptation to urban and rural environments in and around Toronto, Canada in the plant white clover (*Trifolium repens).* White clover exhibits adaptive urban-rural dines in hydrogen cyanide (HCN, an ecologically important plant defense) frequencies in multiple cities around the world (Santan­ gelo et al. 2022b). White clover also displays a multivariate dine across the city of Toronto, whereby numerous quantitative traits (e.g., germination time, flower size) are differentiated along an urbanization gradient (Santangelo et al. 2018). We ad­ dress the following specific questions: (1) Is there evidence of population structure and differentiation among Toronto white clover populations? Addressing this ques­ tion is a prerequisite to disentangling the signatures of selection from the genome­ wide effects of demographic changes and population structure (Hoban et al. 2016). We then ask: (2) Is there evidence of selective sweeps in urban and rural popula­ tions, and if so, which genes show evidence of selection? Our results contribute to our understanding of how the contemporary environmental change in urban and rural habitats influence the evolution of a cosmopolitan plant of economic importance.

## Methods

### Sampling, library preparation, and sequencing

We sequenced a total of 120 plants from 38 populations along three urbanization transects (Figure 1), all of which have been previously associated with phenotypic dines in cyanogenesis (Santangelo et al. 2020b; Thompson et al. 2016), and one of which shows a multivariate cline in multiple quantitative traits (Santangelo et al. 2020a). We sampled 50 plants from 5 urban populations and 45 plants across 21 rural populations, with plants and populations split equally among the three transects (Figure 1). We additionally sequenced 30 plants spread across 12 suburban populations along the three transects. Urban populations were characterized by high levels of the Human Influence Index **(HII),** a composite measure indicating high human population densities and human-associated infrastructure (e.g., built-up areas, artificial light at night, roads, railroads). (Figure 1 inset). While suburban populations ranked just below urban populations in **HII,** rural populations had substantially lower values, consistent with lower levels of human activity and urbanization around these populations (Figure 1 inset).

**Figure 1:**
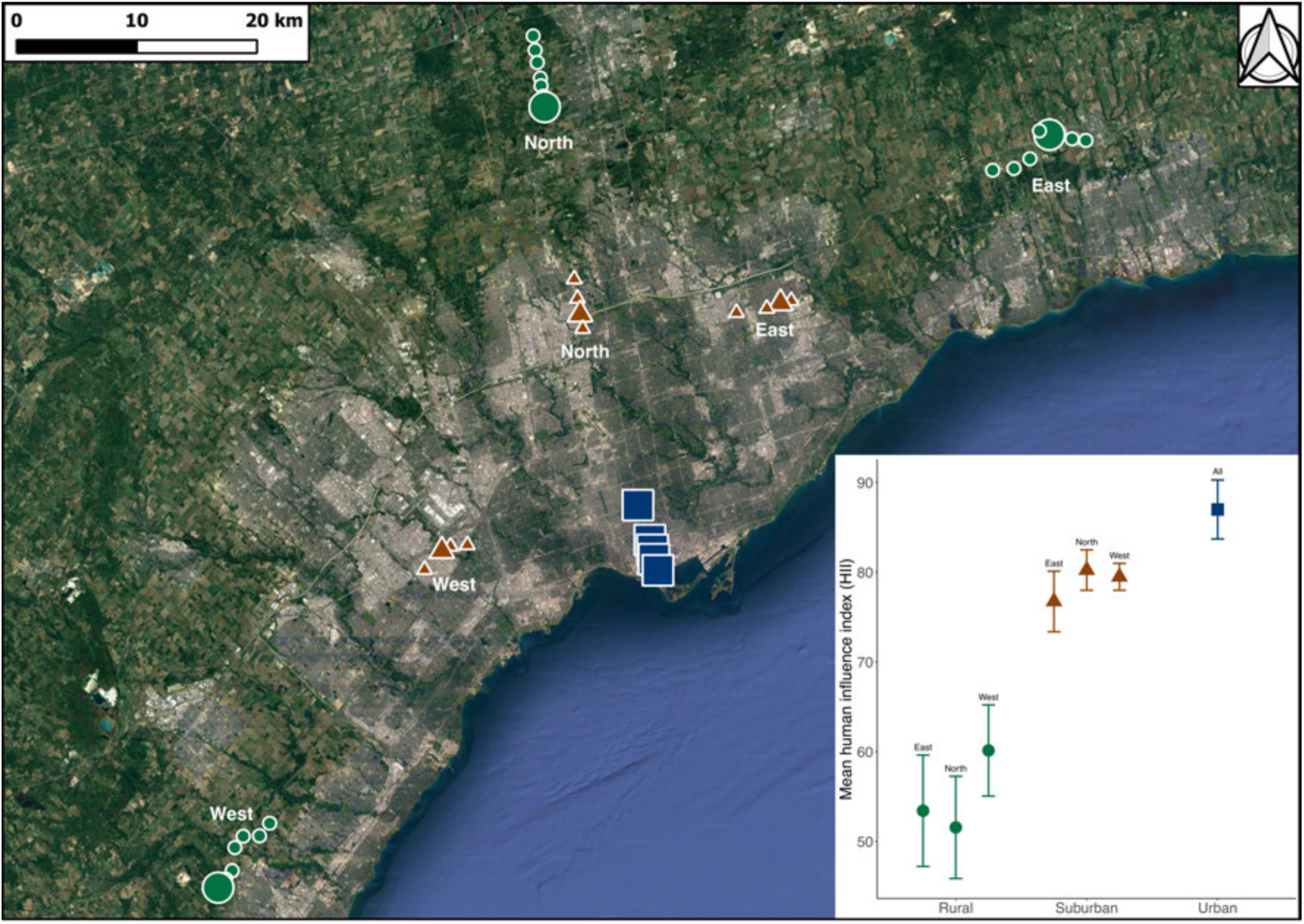
Map of 38 white clover populations sampled for sequencing in Toronto, ON. Sampled populations contained either 1, 7, or 9 samples from rural (green circles), subur­ban (brown triangles), or urban (blue squares) populations. Inset: Mean Human Influence Index **(HII,** ± lSE) of rural, suburban, and urban populations. **HII** for eastern, northern, and western transects are shown separately for rural and suburban populations.

For each plant, we extracted total DNA using a modified CTAB extraction pro­ tocol (Santangelo et al. 2022b) and generated dual-indexed Illumina libraries using an in-house library preparation protocol (Santangelo et al. 2022b). The 120 plants were randomly split into two batches and each batch was sequenced on a single lane of Novaseq S4 using 2 x 150 bp paired-end reads for an expected sequencing depth of ≈13X per sample.

### Sequence alignment and QC

All analyses that follow are available as a publicly-accessible Snakemake pipeline (v.7.21.0, Molder et al. 2021) on GitHub).

We trimmed raw reads using *fastp* v0.23.2 (Chen et al. 2018) with the *-trim_poly_g* argument to trim polyG tails that are commonly added to the ends of reads by No­ vaseq platforms. We performed per-sample QC of both raw and trimmed reads using *FastQC* v0.11.9 (Andrews et al. 2010) and mapped the trimmed reads to the the recently generated *Trifolium repens* haploid mapping assembly (GenBank accession GCA_030408175.1, Santangelo et al. 2023) using *BWA-MEM2* v2.2.1 (Li and Durbin 2009; Vasimuddin et al. 2019). We note that while white clover is a recently-formed allotetraploid (Griffiths et al. 2019), both subgenomes segregate in­ dependently (i.e., inheritance is disomic) and alignments between homeologous loci are rare (Kuo et al. 2024a). This is further supported by high alignment quality scores of short reads across all chromosomes in the present study (except in cen­ tromeres) and within genes on both subgenomes (Figure Sl). Thus, white clover behaves largely as a diploid, both meiotically and bioinformatically (Griffiths et al. 2019; Kuo et al. 2024a; Santangelo et al. 2023).

Following read mapping, we marked duplicate reads and indexed and coordinate­ sorted resulting BAM files using *SAMtools* v1.16.1 (Li et al. 2009) prior to perform­ ing QC of mapped reads using *Qualimap* v2.2.2 (Okonechnikov et al. 2016), *Bam­ tools* v2.5.2 (Barnett et al. 2011), and *multiQC* vl.14 (Ewels et al. 2016). QC of mapped reads revealed three samples with uncharacteristically high alignment error rates and analysis of *rbcL* and *matK* chloroplast genes confirmed that these were *Medicago lupulina.* These samples were removed from all downstream analyses. We removed an additional two samples that had very low sequencing depth (≈1-2X) and unusually high GC% for this species, resulting in 115 high quality samples remain­ ing. Finally, analysis of pairwise relatedness among samples (see below) revealed six sample-pairs with high identity-by-descent *(r*_xy_ > 0.5); we removed one sample from each of these pairs from downstream analyses resulting in a total loss of 11 samples and final sample size of 109 (41 urban, 41 rural, and 27 suburban) with realized sequencing depth of 11.2X.

### Relatedness, population structure, & differentiation

We used genotype likelihoods, which incorporate uncertainty in genotype assign­ ment, for inference of population structure and relatedness to avoid biases associated with calling genotypes in low to medium coverage data (Han et al. 2013; Nielsen et al. 2012, 2011). In addition, we used LD-pruned, 4-fold degenerate sites as putatively neutral sites for these analyses. We began by extracting genome-wide 4-fold sites using *degenotate* vl.2.1 (https://github.com/harvardinformatics/degenotate). We then used *ANGSD* v.0.938 (Korneliussen et al. 2014) to generate Beagle-formatted genotype likelihoods (-doGlf 2) for all 4-fold SNPs (-sites 4-fold.sites, -SNP _pval le-6) using the *SAMtools* genotype likelihood model (-GL 1, Li 2011a) with base alignment quality scores recalculated according to the “extended *SAMtools”* model (-baq 2) to reduce false-positive SNP calls around INDELS (Li 2011b). We only considered reads with a minimum phred-scaled mapping quality of 30 (-minMapQ 30) and sites with a mean base quality of 20 (-minQ 20), minor allele frequency over 0.05 (-minMaf 0.05), and where at least 80% of sequenced individuals (-minind 0.8*N) had at least three reads (-setMinDepthind 3). We additionally ignored sites with more than 2,600X (-setMaxDepth 2600. ≈2 x mean coverage x # samples) to avoid calling SNPs in highly repetitive genomic regions. This resulted in a total of 407,960 4-fold SNPs. We then estimated pairwise LD among these 4-fold SNPs within a 20 kb distance-approximately twice the known breakdown of LD in this system, (Inostraza et al. 2018; Kuo et al. 2024b)-using *ngsLD* v.1.1.1 (Fox et al. 2019) and pruned SNPs using the *prune_graph.pl* script packaged with *ngsLD* al­ lowing a max r^2^ of 0.2 (-min_weight 0.2). This reduced the total number of 4-fold SNPs to 199,297 (≈49%), which were used for inference of population structure and relatedness.

We estimated pairwise relatedness among all 115 high quality samples using *ANGSD* and *NGSrelate* v.2.0 (Hangh0j et al. 2019). We first used *ANGSD* to re­ generate genotype likelihoods at all LD-pruned 4-fold SNPs in binary format (-doGlf 3) using the same filtering criteria as above. We then used *NGSrelate* to estimate pairwise relatedness among all samples, with separate chromosomes processed in parallel. We quantified relatedness using *r_xy,_* which represents the proportion of ho­ mologous alleles shared between two individuals that are identical-by-descent, and is robust to the presence of inbreeding (if present) (Hedrick and Lacy 2015).

We examined patterns of population structure using principle components analy­ sis (PCA) and admixture analyses. We used *PCAngsd* v.0.99 (Meisner and Albrecht­ sen 2018) to estimate the variance-covariance matrix in allele frequencies using the LD-pruned 4-fold SNPs (MAF > 0.05) as input. We then used this matrix as in­ put to the *princomp* function in R v.4.1.3 (R Core Team 2020) to perform a PCA. We performed admixture analyses analyses using *NGSAdmix* v0.933 (Skotte et al. 2013). Using the LD-pruned 4-fold SNPs as input, we ran *NGSAdmix* with ten different random seeds for each assumed number of ancestral clusters (K) between 2 and 10. We then used Evanno’s ΔK method (Evanno et al. 2005) implemented in CLUMPAK v.1.1 (Kopelman et al. 2015) to identify the uppermost level of population structure.

We estimated pairwise nucleotide diversity (*θπ)* within each habitat (urban, rural, suburban) and pairwise differentiation *(F_ST_)* between habitat pairs to complement analyses of population structure. We additionally estimated pairwise population *F_ST_* for all populations containing more than one individual (Figure 1). One- and two-dimensional folded SFS were estimated in *ANGSD.* We first generated separate habitat or population Site Allele Frequency (SAF) likelihood files (-doSaf 1) that were then used to infer lD and 2D site frequency spectra (SFS). We used the same filtering criteria as for the estimation of genotype likelihoods, but with the following modifications: (1) We only required 60% of samples at a site to be covered by one read (-minlnd 0.6 x N, -setMinDepthInd 1); (2) We did not impose a MAF cutoff to avoid biasing the SFS; and (3) We polarized alleles by forcing the reference base to be the major allele in each habitat (-doMajorMinor 4 with -ref) to ensure the same major and minor allele calls in each habitat for *F_ST_* estimates. We then used the *realSFS* script that is packaged with *ANGSD* to estimate folded (-fold 1) one-dimensional urban, suburban, and rural SFS, which were used to estimate per-habitat pairwise nucleotide diversity (*θ_π_),* and two-dimensional habitat (e.g., urban-rural) or population joint SFS, which were used for estimating Hudson’s *F_ST_*(realSFS *F_ST_* index -which *F_ST_* 1, Bhatia et al. 2013; Hudson et al. 1992).

### Signatures of selective sweeps

We initially focused on comparing selection between urban and rural populations because they represented the extremes of our environmental comparisons (Lotterhos and Whitlock 2015), though we note that urban sweep signatures were largely shared with suburban populations (Text S1, Figure S2). In addition, we pooled samples by habitat when detecting signatures of selective sweeps. While one urban and one rural population showed evidence of population structure (populations 40 and 7, respectively, Figure 2), our results do not change if these populations are excluded (Text S2, Figure S3). Finally, it is worth noting that, with the exception of *F_ST_,* the cross-population approaches described below are designed for detecting signatures of positive selection in either urban or rural habitats and are thus not meant to detect regions of divergent selection where alternative alleles in the same region are under selection in urban and rural populations.

**Figure 2:**
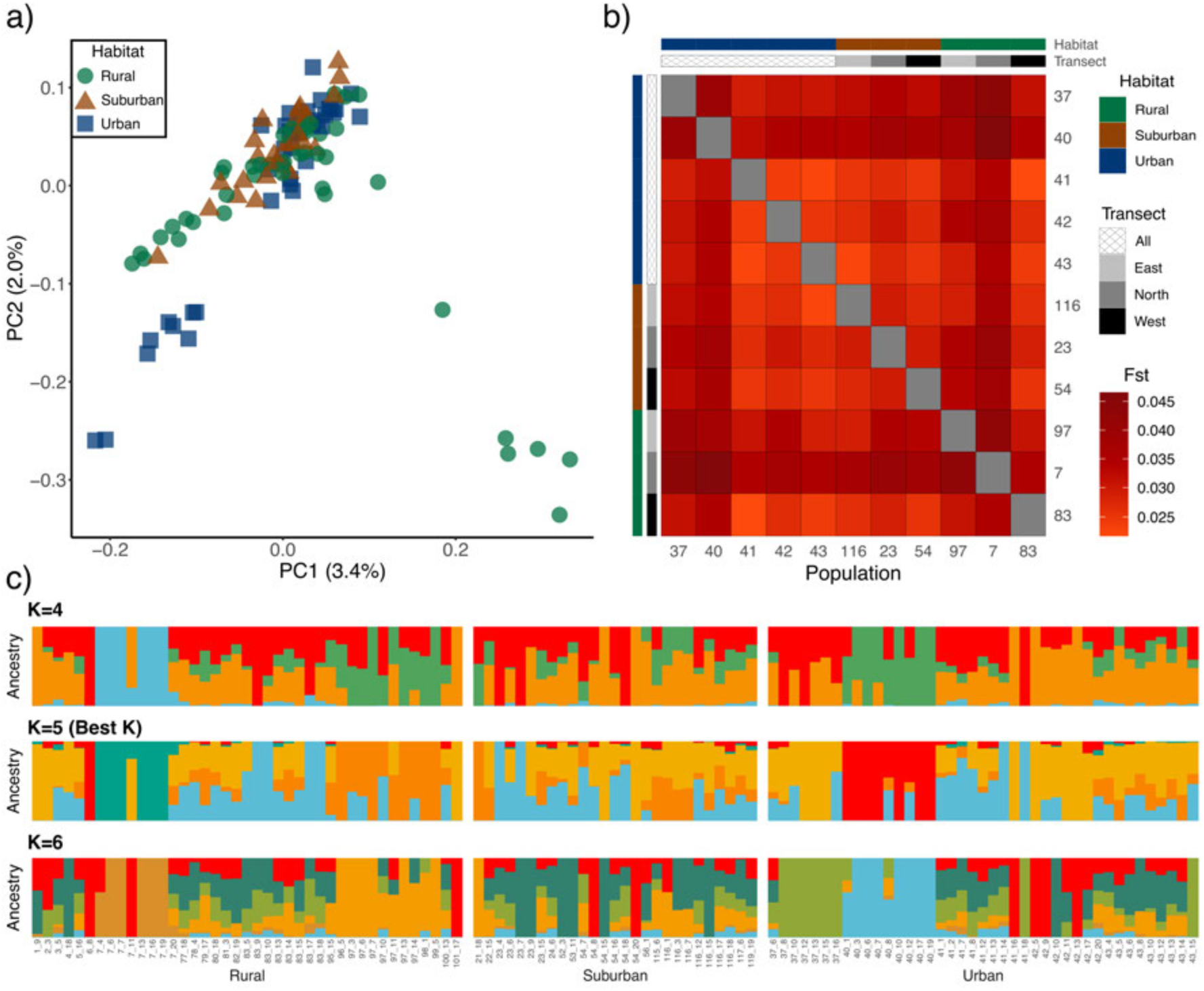
Population structure and admixture among Toronto white clover samples. (a) Principal components analysis (PCA), and (b) Pairwise Hudson’s *F_ST_* between all popula­ tions containing more than 1 individual (N = 11). (c) Admixture proportions for Toronto while clover samples for **K** = 4, **K** = 5 (best **K),** and **K** = 6 assumed ancestral clusters. Individuals are arranged in ascending order by population ID.

To identify urban and rural signatures of selective sweeps, we used approaches based on extended haplotype homozygosity (XP-nSL, nSL, iHS, and iHH12), as well as those based on distortions in the site frequency spectrum (i.e., *F_ST_,* Tajima’s D). We began by calling variants with *freebayes* v.1.3.6 (Garrison and Marth 2012), ignoring regions with greater than 15,000X total coverage (-skip-coverage 15000) to reduce RAM usage. We then filtered VCFs using *bcftools* v.1.16 (Danecek et al. 2021) to include only biallelic SNPs with no missing data, phred-scaled quality scores over 30 *(QU AL>* 30), and additionally removed sites where the number (allele bal­ ance, *AB>=* 0.25 & *AB<=* 0.75 I *AB<=* 0.01), mapping quality *(MQM* >= 30 & *MQMR* >= 30), and paired status *[(PAIRED>* 0.05 & *PAIREDR* > 0.05 & 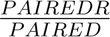 < 1.75 & 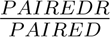 > 0.25) I *(PAI RED<* 0.05 & *PAI REDR* < 0.05)] of reads supporting the reference and alternate alleles were substantially different. Fi­ nally, we removed sites with over 2,600X coverage (i.e., *DP>* 2600) to be consistent with the above population structure analyses and avoid SNPs in highly repetitive regions. Post filtering, 3,917,534 biallelic SNPs remained for downstream analyses. Following filtering, we phased SNPs for downstream haplotype-based scans of selective sweeps. We first performed read-backed phasing using *WhatsHap* vl.7 (Martin et al. 2016) with default parameters. We then used these phase sets with SHAPEIT4 v4.2.2 (Delaneau et al. 2019) to perform population level phasing (-use­ PS 0.0001), which additionally used an interpolated recombination map as input. The recombination map was generated by mapping markers from two recently gener­ ated linkage maps (Olsen et al. 2021) to the white clover reference genome, removing any markers that mapped to the wrong chromosomes, and fitting shape-constrained cubic p-splines with 20 knots to the chromosomal Marey maps (Figure S4). Models were fit using the *scam* function in the *SCAM* vl.2 (Pya and Wood 2015) R package. These models were then used to interpolate the genetic distance for each biallelic SNP retained post-filtering.

We used *selscan* vl.3.0 (Szpiech and Hernandez 2014) to estimate urban-rural cross-population extended haplotype by length (XP-nSL), a haplotype-based statis­ tic with improved power to detect partial and incomplete hard and soft selective sweeps, and which is more robust to genome-wide recombination rate variation than previous approaches (e.g., XP-EHH) (Szpiech et al. 2021). We used the rural habitat as the “reference” population, and the urban habitat used as the “object” popula­ tion. As such, positive XP-nSL scores reflect positive selection occurring in the urban habitat, whereas negative scores reflect positive selection in the rural habi­ tat. After estimating per-site XP-nSL scores, we used *norm* vl.3.0 that is packaged with *selscan* to normalize genome-wide XP-nSL scores. In 50 kb non-overlapping windows, we then estimated the mean normalized XP-nSL score and the proportion of normalized scores> 2 (or< -2). Outlier windows for positive selection in urban habitats were defined as those in the top 1% of the genome-wide empirical mean XP-nSL distribution, and the top 1% of the distribution summarizing the proportion of scores> 2 (Szpiech et al. 2021). Similarly, we defined outlier windows for positive selection in rural habitats as those in the bottom 1% of the genome-wide empirical mean XP-nSL distribution (i.e. the most negative scores), and the top 1% of the distribution summarizing the proportion of scores < -2. To avoid spurious signals driven by noisy XP-nSL estimates in regions with few sampled SNPs, we excluded windows with fewer than 50 SNPs (14.1% of windows, Figure S5). We complemented the XP-nSL analysis by additionally estimating single-population urban and rural segregating sites by length (nSL, Ferrer-Admetlla et al. 2014), integrated haplotype score (iHS, Voight et al. 2006), and iHH12 (Torres et al. 2018), an integrated ver­ sion of the H12 statistic that combines the two most frequent haplotypes and has improved power for detecting soft sweeps (Garud et al. 2015).

Using the tails of empirical distributions as a way of identifying the genomic regions that have experienced selection is common in the literature but has received some criticism, in part, because there is no guarantee that selected loci will occur in the upper tail of empirical distributions, or that loci in the upper tail have experi­ enced selection (Johri et al. 2022). A complimentary approach would be to simulate a null distribution of XP-nSL under the most likely demographic model for our samples and compare our observed values to this null. However, the near absence of population structure between urban and rural white clover populations makes identifying such a model challenging. Instead, we leverage this absence of structure and randomly permuted urban and rural samples 1,000 times, each time estimating XP-nSL in 50 kb windows across the genome, and compared our observed windowed XP-nSL values to this permuted null. We confirmed that the correlation between observed and permuted windowed mean XP-nSL scores were centered around zero across all 1000 permutations (Figure S6). For both the mean windowed XP-nSL score and the proportion of outlier scores within a window, the observed values were in the upper tail of the permuted null, providing further evidence that these are likely candidates of selection (Figure S7).

In addition to approaches based on extended haplotype homozygosity, we leveraged the fact that positive selection creates distortions in the site and frequency spectrum (i.e., SFS). We detected distortions in the SFS by estimating urban-rural *F_ST_* in the same 50 kb windows as our XP-nSL analyses, and considered windows in the top 1% of the genome-wide empirical *F_ST_* distribution as putative candidates of selection. *F_ST_* was estimated using the filtered VCF above in *Pixy* v.1.2.8.betal (Korunes and Samuk 2021), which correctly accounts for the presence of invariant sites and missing data in the calculation of *F_ST_-* We similarly calculated Tajima’s D in urban and rural habitats and estimated the rural-urban difference in Tajima’s D (i.e., *Tajima_Rural_ -Tajima_Urban_)-* We expect that recent selective sweeps would generate more negative Tajima’s D values in the selected habitat in regions of the genome that are putative candidates of selection.

### Clinal change in allele frequencies using suburban popula­ tions

We were interested in examining whether regions with the strongest evidence of selective sweeps in urban or rural habitats showed any evidence of clinal change in allele frequencies. To do this, we identified the site with the maximum urban-rural *F_ST_* within each of our top candidate regions for selection in urban and rural habitats (Table 1), and estimated the minor allele frequency at this site in each of our populations that contained more than one individual (Figure 1). For each of these sites, we ran a binomial regression with minor allele frequency as the response variable and distance from the urban core (in Km) as the sole predictor, consistent with previous work that examined clinal change in cyanogenesis frequency along urban­ rural gradients (Santangelo et al. 2022b). In addition, we compared the urban-rural difference in minor allele frequency to the suburban-rural and suburban-urban difference to understand whether allele frequencies of suburban populations were more similar to those of urban or rural populations at these most differentiated sites. For each of these analyses, we included 100 randomly selected sites from across the genome for comparison.

**Table 1:** Regions showing the strongest evidence of positive selection in urban and rural habitats (i.e., XP-nSL outliers additionally in the top 5% of the urban-rural permuted XP-nSL distributions). For each region, we show it’s position (Chromosome, Start, End); size (as a multiple of 50 kb where consecutive outlier windows were merged); the habitat under selection; the mean XP-nSL score; the minimum (for rural) or maximum (for urban) XP-nSL score· the proportion of raw scores greater than 2 (for urban) or less than -2 (for rural) for each window; the percentile of the urban-rural permuted null for mean XP-nSL scores in which the windows lie; the percentile of the urban-rural permuted null for the proportion of scores exceeding the threshold in which the windows lie; the rank of each outlier window in the concatenated region; mean Tajima s D across the region in urban and rural habitats· any named annotated genes in the region; and the figures corresponding to that region. For regions containing multiple outlier windows, proportions, percentiles, and ranks report values for each window. Ranks are based on the “Percentile of proportion permuted XP-nSL distribution’ column.

| Chromosome | Start | End | Region size | Selection | Mean XP-nSL | Min/Max XP-nSL | Proportion XP-nSL outliers | Percentile of mean permuted XP-nSL distribution | Percentile of proportion permuted XP-nSL distribution | XP-nSL ranks | Tajima’s D | Annotated genes | Figure |
| --- | --- | --- | --- | --- | --- | --- | --- | --- | --- | --- | --- | --- | --- |
| 16 | 16600000 | 16700000 | 100000 | Rural | -4.33 | -8.09 | 1,0.825 | 1,0.991 | 1,0.983 | 1,5 | Urban = -0.55<br>Rural = -1.09 | ITPK1, SUVH2, CML18, SYP22, ABCB2 | Figure 4d, S13 |
| 13 | 49750000 | 49900000 | 150000 | Rural | -2.82 | -6.26 | 0.774,0.949,0.66 | 0.951,0.997,0.947 | 0.974,0.996,0.939 | 7,2,18 | Urban = 0.14<br>Rural = -0.74 | MCM8, RPS23B, RH57, NFYB8, RPL32A |  |
| 9 | 56600000 | 56650000 | 50000 | Rural | -2.89 | -4.35 | 0.901 | 0.985 | 0.992 | 3 | Urban = -0.34<br>Rural = -0.81 | ORC4 |  |
| 10 | 58150000 | 58200000 | 50000 | Rural | -2.85 | -4.65 | 0.843 | 0.983 | 0.985 | 4 |  | ABCG26, CBL3, HSC80 |  |
| 7 | 7350000 | 7600000 | 250000 | Rural | -2.36 | -5.45 | 0.791,0.743,0.538, 0.749,0.632 | 0.959,0.937,0.908, 0.971,0.894 | 0.977,0.967,0.849, 0.968,0.925 | 6,9,30,8,20 |  | ACA11, GPDHC1, CKX6, TIP51, MAN2, HSP22, MEX1, IQD11, UBP24, COG7, TOP1A | Figure 4a, S10 |
| 14 | 49450000 | 49500000 | 50000 | Rural | -3.03 | -5.81 | 0.727 | 0.989 | 0.963 | 11 |  | RH57 |  |
| 9 | 16400000 | 16450000 | 50000 | Rural | -2.74 | -5.42 | 0.711 | 0.978 | 0.958 | 14 |  | IWS1 |  |
| 10 | 58250000 | 58350000 | 100000 | Rural | -2.51 | -5.39 | 0.563,0.706 | 0.955,0.965 | 0.874,0.956 | 29,15 | Urban = -0.12<br>Rural = -0.92 | UKL4, SMC3, RFS1 |  |
| 9 | 4350000 | 4700000 | 350000 | Urban | 2.32 | 4.41 | 0.4,0.442,0.682,1, 0.714,0.527,0.841 | 0.787,0.734,0.932,0.989, 0.968,0.872,0.962 | 0.568,0.684,0.949,1, 0.96,0.835,0.986 | 77,59,17,1, 12,38,7 | Urban = -1.60<br>Rural = -1.08 | AP2M, ALMT12, ABCI7, PER20, TOM2A, APRR2, RABH1B, LON2, UPF1 | Figure 4c, S12 |
| 9 | 4800000 | 4900000 | 100000 | Urban | 3.50 | 6.67 | 0.95,0.917 | 1,0.962 | 0.996,0.993 | 2,4 | Urban = -0.67<br>Rural = -0.32 | PALM1 | Figure 4c, S12 |
| 7 | 51000000 | 51100000 | 100000 | Urban | 2.43 | 4.73 | 0.935,0.481 | 0.979,0.89 | 0.994,0.765 | 3,48 | Urban = -0.62<br>Rural = -0.23 | FKBP13 | Figure 4b, S11 |
| 7 | 53600000 | 53900000 | 300000 | Urban | 2.70 | 6.22 | 0.636,0.904,0.636, 0.761,0.521,0.796 | 0.89,0.995,0.917, 0.995,0.901,0.99 | 0.928,0.992,0.928, 0.972,0.828,0.979 | 21,5,20, 10,40,9 | Urban = -0.61<br>Rural = -0.11 | AKHSDH1, AAE15, SHM4 | Figure 4b, S11 |
| 5 | 56900000 | 57000000 | 100000 | Urban | 2.47 | 4.76 | 0.553,0.862 | 0.885,0.983 | 0.865,0.988 | 30,6 | Urban = -1.10<br>Rural = -0.78 | TUB5 |  |
| 16 | 350000 | 450000 | 100000 | Urban | 2.27 | 3.83 | 0.606,0.797 | 0.862,0.959 | 0.91,0.98 | 25,8 |  | DJA6 |  |
| 5 | 46150000 | 46200000 | 50000 | Urban | 2.47 | 4.78 | 0.709 | 0.957 | 0.958 | 13 |  |  |  |
| 5 | 32400000 | 32450000 | 50000 | Urban | 2.46 | 4.36 | 0.695 | 0.956 | 0.953 | 15 |  |  |  |

### Functional characterization of selected regions

To examine candidate genes that may be targets of positive selection in urban and rural populations, we intersected our urban and rural XP-nSL outlier windows with the annotation file (GFF3) accompanying the recently published white clover reference genome (Santangelo et al. 2023). We focus only on regions that fall in the top 5% of the urban-rural permuted XP-nSL distribution. In addition, we performed a Gene Ontology analysis for Biological Processes and Molecular Functions separately for urban and rural selected regions using the *topGO* v.2.50.0 (Alexa and Rahnenfuhrer 2016) R package with Fisher’s exact test, the “weight01” algorithm and a *P-value* cutoff of 0.05. We did not correct for multiple testing since the resulting corrected p-values are often overly conservative, and by accounting for the hierarchical structure of the GO topology, the p-values estimated by the “weightOl” algorithm are not independent, violating a key assumption of multiple testing correction (Alexa and Rahnenfuhrer 2016).

## Results and Discussion

### Genetic diversity, differentiation, and population structure

Urban, rural, and suburban habitats showed identical levels of pairwise nucleotide diversity at four-fold degenerate sites *_(πRural_* = *_πSuburban =πUrban=_* 0.0187). There was little evidence of population structure from the PCA, whereby the first two PCs together accounted for ≈5.4% of genetic variation across samples (PC1 = 3.4%, PC2 = **2%)** and, with the exception of one rural and one urban population, showed no evidence of clustering by habitat (Figure 2a). Admixture analyses and Evanno’s **ΔK** method identified **K** = 5 as the most likely number of ancestral genetic clusters, and with the exception of the same urban and rural population as in the above **PCA,** all clusters were well represented in each habitat regardless of the value of K (Figure 2c). The low levels of population structure among Toronto white clover samples were further supported by low levels of genetic differentiation at four-fold sites between habitats (Rural-Suburban *F_ST_* = 0.0060, Rural-Urban *F_ST_* = 0.0063, Urban-Suburban *F_ST_* = 0.0062) and population pairs (0.02 <= *F_ST_* <= 0.046, Figure 2b).

Spatially-restricted gene flow combined with local genetic drift can lead to rapid population differentiation and strong patterns of population structure. Indeed, habi­ tat fragmentation and reductions in population size associated with urbanization are frequently cited drivers of reduced genetic diversity in urban populations and/or in­ creased urban-rural genetic differentiation in rodents (Combs et al. 2018; Harris et al. 2016; Munshi-South 2012; Munshi-South and Kharchenko 2010), lizards (Beninde et al. 2018, 2016; Littleford-Colquhoun et al. 2017; Lourenço et al. 2017; Noel et al. 2007), birds (Bjorklund et al. 2010; Mueller et al. 2018; Unfried et al. 2013), mam­ mals writ-large (Schmidt et al. 2020), and plants (Bartlewicz et al. 2015; Rivkin and Johnson 2022; Yakub and Tiffin 2017). However, a recent meta-analysis found only weak effects of urbanization on within-population genetic diversity and no consistent effect on between-population differentiation, and instead suggested that patterns are largely system-dependent because not all species are equally affected by urban fragmentation (Miles et al. 2019). Toronto white clover appears to be a particularly extreme case, with high genetic diversity in all habitats, low genetic differentiation between habitats, and high levels of admixture that together lead to overall weak urban-rural population structure (Figure 2) and an absence of isolation-by-distance at this scale (Mantel’s r = 0.13, *P* = 0.26, Figure S8). These results are consistent with previous work showing generally low levels of differentiation and urban-rural structure in this system in multiple cities around the world (Caizergues et al. 2024; Johnson et al. 2018; Santangelo et al. 2022b). White clover is a perennial, obligate outcrosser that maintains large populations, and is likely repeatedly introduced into cities from multiple sources due its use in agriculture and presence in turf mix­ tures (KjIBrgaard 2003; Sincik and Acikgoz 2007). In addition, while gene flow via pollinator-mediated pollen dispersal likely occurs within a limited geographical range in this system (De Lucas et al. 2012; L0jtnant et al. 2012), white clover seeds are among the most commonly detected species found on human clothing, vehicles, and animals (Pickering and Mount 2010). Therefore, longer-distance human- or animal-mediated seed dispersal may be particularly important in this system, and this is likely more pronounced in and around cities where the dispersal agents (e.g., humans) are more common. In short, white clover is unlikely to have its population sizes reduced by urban fragmentation, and instead likely maintains high population connectivity and diversity in Toronto and other cities around the world.

### Signatures of selection in urban and rural populations

We identified 130 50 kb XP-nSL outlier windows with evidence of positive selec­ tion in urban habitats, and 123 windows with evidence of positive selection in rural habitats (Figure 3), which reduce to 102 and 105 outlier regions in urban and ru­ ral populations after merging consecutive outlier windows, respectively. However, because outlier approaches can be enriched for false positive signals of selection, we focus our attention on outlier windows that are additionally in the upper 5% of the urban-rural permuted XP-nSL distribution (diamonds in Figure 3b, Figure S7), representing 13 urban and 11 rural outlier windows combined into 8 urban and 8 rural outlier regions (Table 1). We additionally note that many of these regions ex­ hibit more negative Tajima’s D in the habitat that is the putative target of positive selection, consistent with the accumulation of new variation after a recent selective sweep (Figure 3b, Table 1).

**Figure 3:**
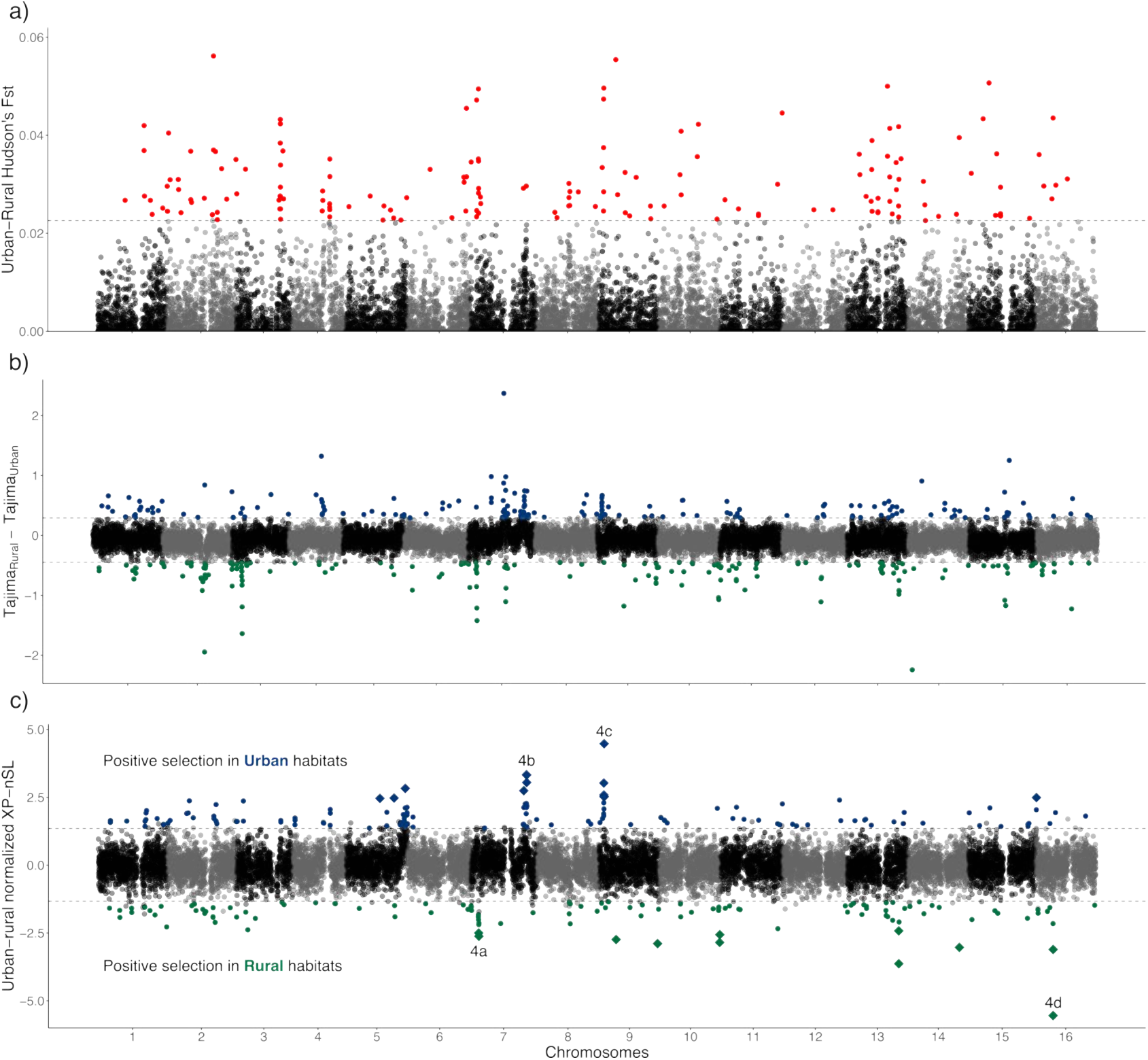
(a) Urban rural Hudson’s *F_ST_* estimated in 50 kb windows across the genome. Red points indicate windows in the top 1% of the genome-wide empirical distribution. (b) Difference between rural and urban Tajima’s D (i.e., *Tajima_Rural_*-*Tajima_Urban_* estimated in 50 kb windows across the genome. Blue and green points are in the top and bottom 1% of the genome-wide empirical *Tajima_Rural_*-*Tajima_Urban_* distribution, respectively. Blue points are consistent with selective sweeps in urban habitats (i.e., *Tajima_Urban_*< *Tajima_Ruraz_),* while green points are consistent with sweeps in rural habi­ tats (i.e., *Tajima_Rural_* < *Tajima_Urban_)* (c) Urban-rural normalized XP-nSL estimated in 50 kb windows across the genome. Blue points represent outliers showing evidence of pos­ itive selection in urban habitats, while green points represent windows showing evidence of positive selection in rural habitats. Diamonds represent windows that are additionally in the top 5% of the urban-rural permuted XP-nSL distribution (Figure S7). Letters in (c) correspond to panels (a)-(d) in Figure 4, where magnified Manhattan plots are shown for those Figure 4: Magnified Manhattan plots showing per-site normalized urban-rural XP-nSL scores for regions specified in Figure 3c. Coloured horizontal lines at the bottom of each panel show regions that are also in the top 1% of the genome-wide distribution for other SFS- and Haplotype-based statistics: *F_ST_*(red), nSL (orange), iHS (purple), and **iHH12** (green). Light grey bands display XP-nSL outlier windows, while dark grey bands are XP­ nSL windows that are additionally in the top 5% of the urban-rural permuted XP-nSL distribution. Yellow diamonds in (c) and (d) show the maximum and minimum genome­ wide XP-nSL scores, respectively.

Among the strongest candidates of positive selection in urban populations is a 550 kb region on chromosome 9 (Figure 4c, Figure S9). This region contained three genome-wide XP-nSL outlier regions, two of which were additionally outliers in the urban-rural permuted XP-nSL distribution (Table 1). In addition, these regions are overlapped by five *F_ST_* outlier windows, and six nSL, two iHS, and six iHH12 urban outlier windows (Figure 4c), and contain the maximum genome­ wide XP-nSL score (yellow diamond in Figure 4c). Similarly, an approximately 2.9 Mb region on chromosome 7 (Figure 4b, Figure SlO) contained seven outlier regions, four of which were in the top 5% of the urban-rural permuted XP-nSL distribution (Table 1). This region was additionally overlapped by two *F_ST_* outliers, and contain three nSL outlier windows. These regions on chromosomes 7 and 9 contain 44 and 48 genes, respectively, which are collectively involved in biotic and abiotic stress, such as plant pathogen immunity (*TOM2A,* Tsujimoto et al. 2003), salt stress tolerance *(APRR2,* Jiang et al. 2022), and drought/cold stress *(AAE15,* Shockey and Browse 2011), as well as in growth and metabolism such as seedling germination and cotyledon growth *(RABHJB* and *LON2,* He et al. 2018; Tsitsekian et al. 2019), and leaf formation and development *(PALMJ,* Chen et al. 2010). The highest genome-wide XP-nSL score occurred within the coding sequence of a gene encoding an Acetylornithine deacetylase (NAOD), which plays a critical role in arginine and polyamine biosynthesis and is known to impact flowering time and fruit ripening in *Arabidopsis* (Molesini et al. 2017).

**Figure 4:**
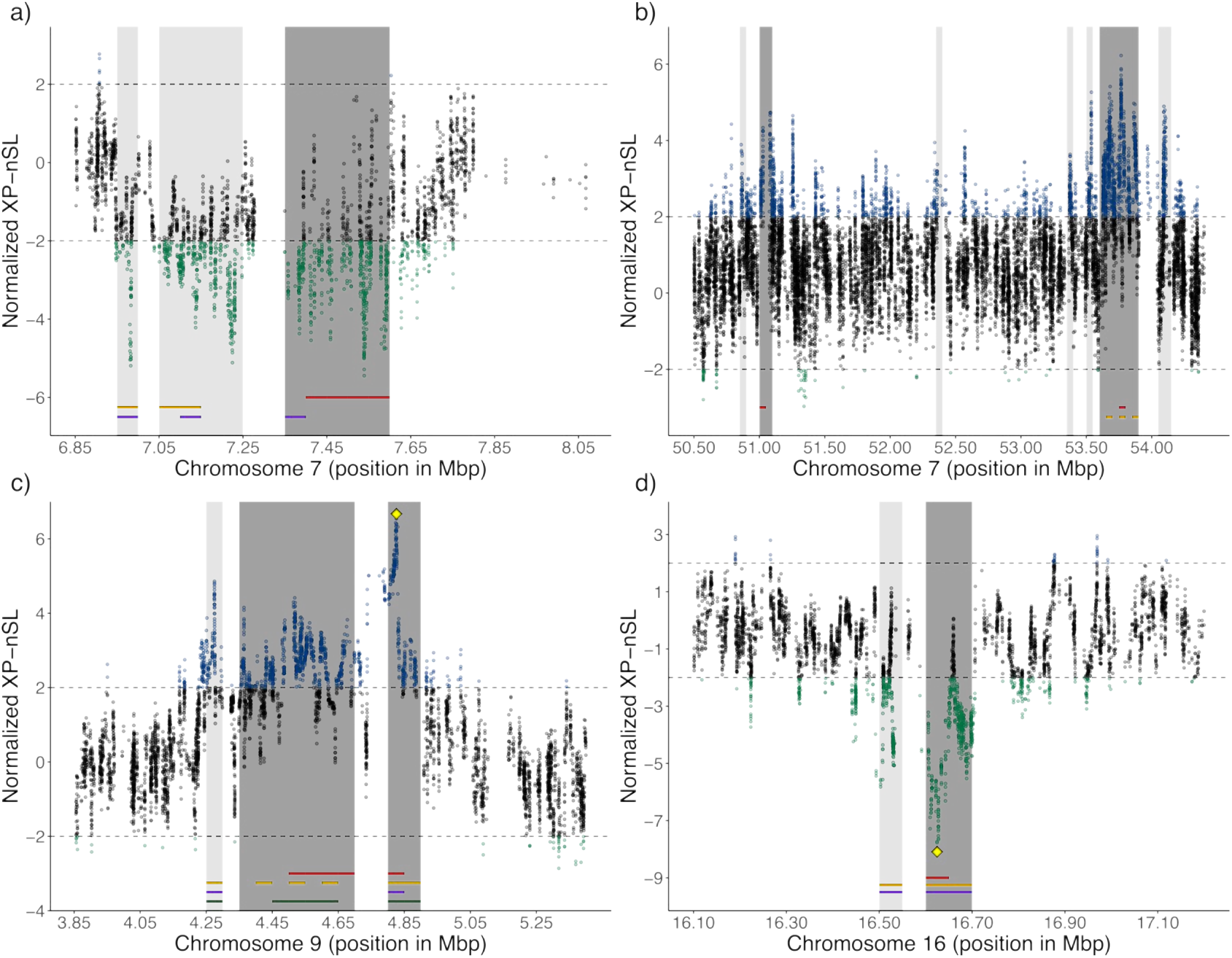
Magnified Manhattan plots showing per-site normalized urban-rural XP-nSL scores for regions specified in Figure 3c. Coloured horizontal lines at the bottom of each panel show regions that are also in the top 1% of the genome-wide distribution for other SFS- and Haplotype-based statistics: *F_ST_* (red), nSL (orange), iHS (purple), and iHH12 (green). Light grey bands display XP-nSL outlier windows, while dark grey bands are XPnSL windows that are additionally in the top 5% of the urban-rural permuted XP-nSL distribution. Yellow diamonds in (c) and (d) show the maximum and minimum genomewide XP-nSL scores, respectively.

Rural populations showed similarly strong signatures of selection across the genome. For example, a 650 kb region on chromosome 7 (Figure 4a, Figure S11) contained three outlier regions with one 250 kb region that was additionally an out­ lier in urban-rural permuted XP-nSL distribution (Table 1). These regions were overlapped by four *F_ST_* outlier windows, and three nSL and iHS rural outlier win­ dows. Similarly, a 200 kb region on chromosome 16 (Figure 4d, Figure S12) contains three outlier regions, with one 100 kb region that is an outlier in the urban-rural permuted XP-nSL distribution (Table 1). This region is overlapped by two *F_ST_* outlier windows and three nSL and iHS rural outlier windows, and contains the genome-wide minimum XP-nSL score (yellow diamond in Figure 4d). These regions on chromosomes 7 and 16 contain 25 and 11 genes, respectively, and are involved in the development of plant structures such as roots (*GPDHCJ* and *ITPK1,* Hu et al. 2014; Pullagurla et al. 2023), as well as resistance to abiotic stresses such as shade (*CKX6,* Wang et al. 2020), heat *(HSP22,* Yamaguchi et al. 2021), frost, *(MEX1,* Cvetkovic et al. 2021), and salt and drought tolerance (*UBP24* and *SUVH2,* Guo et al. 2025; Wu et al. 2019).

Both urban and rural land-use change have threatened biodiversity worldwide and altered the ecological conditions experienced by natural populations (Johnson and Munshi-South 2017; Newbold et al. 2015; Turcotte et al. 2017). Consistent with this view, we identified signatures of positive selection in both habitats in genes underlying tolerance to stressful biotic and abiotic conditions. Similarly, sweep signatures in genes related to plant vegetative (e.g., leaves and roots) and floral growth and development are consistent with previous common-garden work documenting urban-rural differentiation in vegetative biomass, flowering time, germination time, and flower size among plants from many of the same populations as those investigated here (Santangelo et al. 2020a). Gene Ontology (GO) analysis of candidate genes in regions with the strongest evidence of selection (Table 1) support both urban and rural populations experiencing sweeps in regions underlying growth, development, and tolerance to biotic and abiotic stress. However, these are likely occurring through different pathways and mechanisms (Table Sl and Table S2); the inositol signaling pathway-a pathway known to influence tolerance to multiple abiotic stressors in plants (Jia et al. 2019)-seems to be particularly important among plants from rural Toronto white clover populations, while urban populations appear to leverage other means (e.g., UDP-glycosyltransferases, Table Sl, Table S2, Gharabli et al. 2023). Importantly, to confirm these results, future work is required to better characterize the functional relevance of genes and genomic regions underlying adaptation to urban and rural habitats.

White clover has long served as a model for understanding the genetic basis of adaptation due to the presence of temperature-mediated cyanogenesis dines across multiple continents (Daday 1954, 1965; Innes et al. 2022; Kooyers and Olsen 2013), and recent detection of similar dines across urban-rural gradients putatively driven by the same selective agents (Santangelo et al. 2022b). The recent release of multiple high quality reference genomes for the species (Kuo et al. 2024a; Santangelo et al. 2023) has spurred investigations into other genomic regions underlying local adaptation in white clover. For example, recent work has shown that, while cyanogenesis frequencies are correlated with temperature across North America, local adapta­ tion is instead driven primarily by variation in genes underlying growth and flow­ ering in response to seasonal temperature cues (Kuo et al. 2024b). Similarly, white clover’s invasion of the Americas from the Mediterranean seems to be associated with variation around multiple, large, fitness-associated structural variants that af­ fect differential expression of genes underlying growth, defense, and stress tolerance (Battlay et al. 2024). Three of these-two on chromosome 7 and one on chromo­ some 9-form allele frequency dines in North America (haploblocks HB7a2, HB7b, and HB9, Battlay et al. 2024) and perfectly overlap the strongest signals of positive selection reported here in rural (HB7a2: Figure 4a) and urban (HB7b: Figure 4b, HB9: Figure 4c) populations, though we note that only two of these (Hb7b and Hb9) display urban-rural allele frequency differences in Toronto white clover pop­ ulations *(Hb7b_Urban_* = 0.13, *Hb7brur_al_* = 0.23; *Hb9_Urban_* = 0.037, *Hb9_rural_* = 0.160).

In addition to facilitating white clover’s rapid adaptation following invasion, these regions may be similarly important in driving local adaptation along the smaller spatial scale of urban-rural gradients in Toronto. While recombination-suppressing structural variants (e.g., inversions) may generate false positive signals of selection in these regions, these haploblocks do not appear to be associated with genome-wide regions of low recombination (Figure S13), likely because recombination within ho­ mozygotes is still ongoing. Additionally, xp-nSL is strictly comparative and therefore would be capable of indentifying long haplotypes in one population regardless of the inversion. Moreover, Tajima’s D values in Hb7b and Hb9 are negative in the selected populations, consistent with recent positive selection (Table 1). However, neither the polarization into ancestral/ derived nor the phenotypic effects of these structural variants are currently known, and thus future examination of historical sequences (e.g., herbarium specimens) and/or urban-rural reciprocal transplants that link hap­ loblock frequencies to phenotypes and fitness will be important for quantifying the role that these regions play in urban and rural adaptation. In addition, ongomg and future work will investigate the extent to which these sweep signatures are shared across cities, and experimentally testing the agents and targets of selection underlying white clover’s adaptation to urban and rural habitats.

We have focused on characterizing selective sweeps in urban and rural popula­ tions because these represent the extremes of our environmental gradient, and urban sweep signatures are largely shared with suburban populations (Text S1, Figure S2). Nonetheless, it is worth noting that urban and suburban habitats show some evi­ dence of differentially selected regions across the genome (Text S1, Figure S2), and many of the most strongly selected regions (Table 1) appear to show clinal change in allele frequency at the most differentiated SNP (Figure 5, Figure S14). We note that within these regions, suburban allele frequencies were closer on average to urban populations than to rural populations (Figure S14b), which is again consistent with the selective environment of suburban habitats more closely matching that of urban, rather than rural, populations. Nonetheless, while early work in urban evolutionary ecology tended to pool urban and suburban populations together, there has been increasing emphasis on the importance of considering heterogeneity in ecological and evolutionary responses among populations within urban spaces (Des Roches et al. 2020; Martin et al. 2024; Prileson and Martin 2024; Santangelo et al. 2022a), and similar arguments can surely be made of rural habitats. Our results show that, al­ though the strongest differences in genome-wide signatures of selective sweeps occur between urban and rural habitats, there may also be fine-scale adaptation within occurring within populations cities that warrants further consideration.

**Figure 5:**
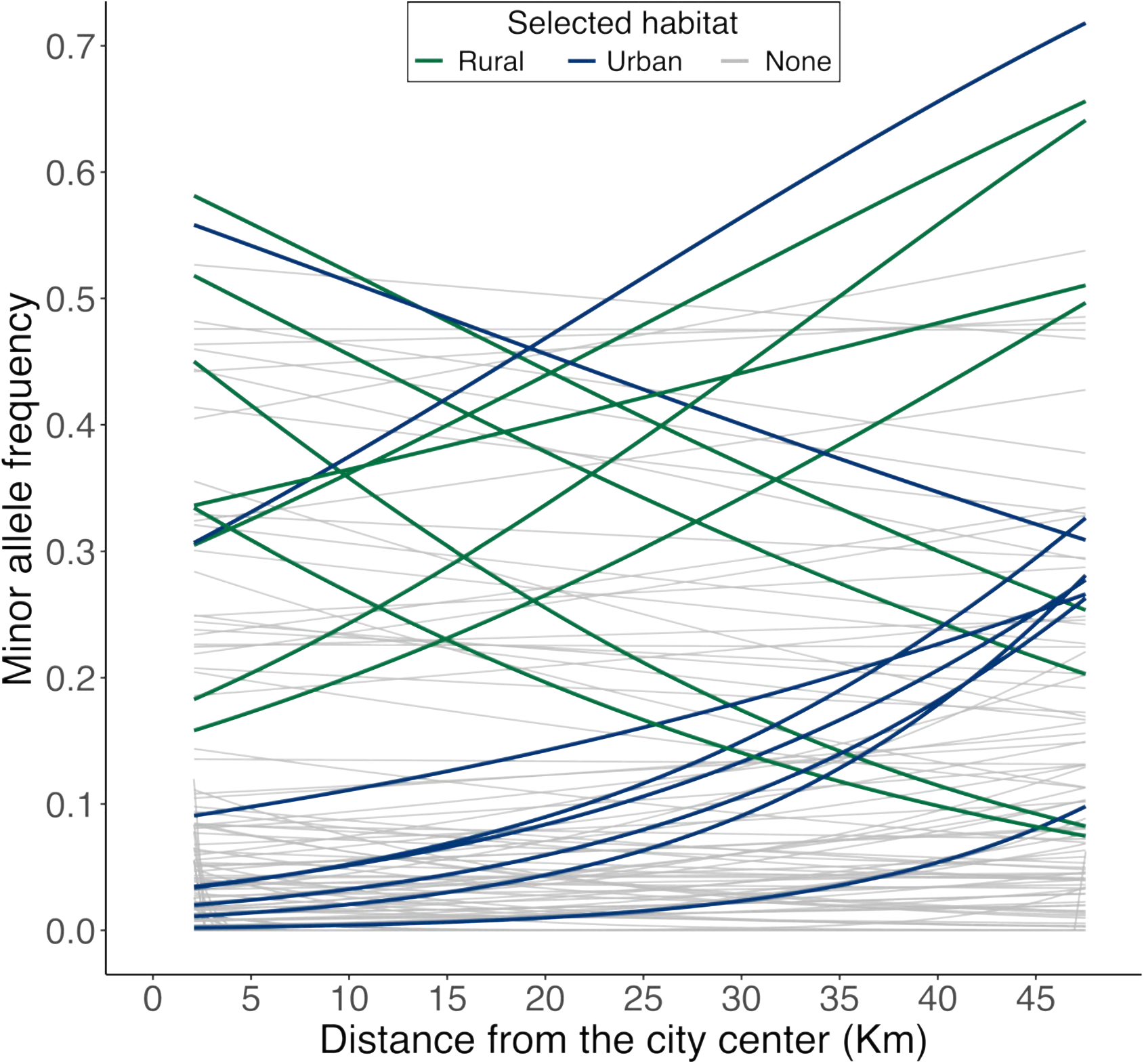
Minor allele frequency clines at the most differentiated SNP in each of the top candidate regions of selection (Table 1). Lines are colored based on the inferred direction of selection from the XP-nSL analysis with blue lines indicating positive selection in urban habitats and green lines reflecting positive selection in rural habitats. Grey lines show allele frequency clines for 100 randomly selected SNPs across the genome. Similar version but showing MAF by habitat rather than distance in shown in Figure S14.

### Characterization of urban and rural sweeps signatures

Local adaptation requires that spatially divergent selection overcome the homoge­ nizing effects of gene flow. For species with high levels of population connectivity and migration, local adaptation is predicted to be driven primarily by loci of large­ effect (Hodgins and Yeaman 2019; Lenormand 2002; Yeaman 2022). This may be especially true in the early phases of adaptation following a rapid shift to a new environment (Hayward and Sella 2022), as may be expected when natural habitat is converted to cities or agricultural land. Our results support this view, as many of our strongest signatures of positive selection in both habitats overlapped known structural variants with large effects on fitness (Battlay et al. 2024). These structural variants are segregating at appreciable frequencies worldwide, suggesting adaptation is occurring via selection on standing genetic variation in these regions, which may be expected given Toronto’s relatively young age. However, we note that *F_ST_* values between urban and rural populations in Toronto in these regions are generally low and that these regions likely represent incomplete sweeps with ongoing selection, which are better detected by haplotype-based statistics (e.g., XP-nSL) than *F_ST_* (Szpiech et al. 2021). Patterns of allele frequency and haplotype differentiation in these regions seem to support this, whereby haplotypes in the most strongly selected regions only weakly cluster by habitat (Figure S9-Figure S12), though clustering is still substantially greater than unselected regions of the genome (Figure S15). An alternative, though not mutually exclusive, reason for the generally low *F_ST_* values could be that selection is highly polygenic with diffuse signals spread across the genome, driven by selection across multiple traits and along multiple biotic and abiotic axes in urban and rural habitats. In support of this, we found that chro­ mosomes with more genes contained more **XP-nSL** outlier loci in both urban and rural habitats (Figure S16). This result is consistent with recent work in great tits that found polygenic signals of urban adaptation (Salmon et al. 2021). Ongoing and future work leveraging sequence data from phenotyped plants will more thoroughly examine the relative roles of large-effect loci and diffuse polygenic architectures to adaptation in urban and rural habitats.

The relatively young age of cities like Toronto suggests that the bulk of adapta­ tion is likely to be driven by selection acting on standing genetic variation. However, the origin of this variation is particularly interesting in a system like white clover given that it is still routinely cultivated for its use in agriculture. Indeed, white clover cultivars have historically been bred for increased persistence and often tar­ geting traits involved in resistance to abiotic (e.g., drought, frost) and biotic (e.g., disease, herbivory) stress, and morphological and life-history traits such as flower­ ing intensity, stolon number, seed yield, and biomass productions (Taylor 2008). Many of these traits may be similarly adaptive in naturalized urban or rural pop­ ulations. While Canada has historically used a only a limited number of cultivars in agricultural settings (e.g., “white Dutch” from Europe or “wild white” from the United Kingdom or New Zealand, Canada. Dept. of Agriculture. 1941), seed from many common cultivars can be purchased online and are increasingly planted in gar­ dens as “clover lawns”. Therefore, adaptation of urban and rural white clover may represent a mosaic of selection acting on variation from natural populations and in­ trogressed variation from commercially produced cultivars. Identifying the cultivars most frequently planted around our study populations coupled with whole-genome sequencing and phenotyping of these cultivars will help shed light on the potential role of introgression in driving urban and rural adaptation of Toronto white clover populations.

## Conclusion

Urbanization results in rapid and drastic environmental changes with profound con­ sequences for the ecology and evolution of natural populations. Urban fragmenta­ tion frequently drives reduced population size and connectivity, leading to reduced diversity and increased differentiation and population structure for many species. However, white clover appears largely unaffected by urban fragmentation, instead maintaining high diversity across all habitats with little detectable structure be­ tween urban and rural populations, likely resulting from high dispersal across the city in this obligately outcrossing perennial. While much focus has been placed on phenotypic and genomic responses to urbanization, rural habitats can similarly im­ pose strong anthropogenic selection. Indeed we find signatures of selective sweeps in regions underlying growth, reproduction, and biotic and abiotic stress tolerance in both urban and rural Toronto white clover populations, and SNPs in these regions exhibit clinal variation in allele frequencies across our three transects. Preliminary characterization of selective sweeps suggests that selection is likely ongoing (i.e., incomplete sweeps) and, while some of our strongest signals of selective sweeps over­ lap known large-effect structural variants, we detect a general signal of polygenic adaptation in both urban and rural habitats. As increased functional genomic in­ formation becomes available in this system, will will be better positioned for a more thorough characterization of the genetic basis and architecture of adaptation to these contemporary anthropogenic environments.

## Data Accessibility

All code used to produce the manuscript’s results can be found on the GitHub page for JSS (https://github.com/James-S-Santangelo/toronto_gwsd) and ad­ ditionally archived on Zenodo (will provide final DOI upon acceptance). Raw reads for all samples have been deposited on GenBank (BioProject PRJNAll 79961).

## Author Contributions

**JSS:** Conceptualization, Methodology, Software, Formal analysis, Data curation, Writing - Original draft, Writing - Review & Editing, Visualization, Project admin­ istration **MTJJ:** Conceptualization, Methodology, Resources, Writing - Review & Editing, Visualization, Supervision, Funding Acquisition **RWN:** Conceptualization, Methodology, Resources, Writing - Review & Editing, Visualization, Supervision, Funding Acquisition

## Acknowledgments

We would like to thank Ken A. Thompson for the initial plant collections from which the genomic data in this manuscript were produced. We would also like to thank the computational resources provided by the Digital Research Alliance of Canada.

## Supplementary Text

### Text S1: Sweep signatures involving suburban populations

In addition to testing for signatures of positive selection between urban and ru­ ral populations, we used our phased haplotype data to compare compare selective sweep signatures between suburban and rural populations, and between urban and suburban populations. Specifically, we used *selscan* with suburban populations as the “object” population and rural populations as the “reference” (Figure Sla), and again with urban populations as the “object” and suburban populations as the “reference” (Figure Slb). Of the 135 genome-wide windows with evidence of pos­ itive selection in urban populations from the urban-rural comparison (Figure 3b), 15.5% were shared with suburban populations (Figure Sla). By contrast, of the 124 genome-wide windows with evidence of positive selection in rural populations from the urban-rural comparison (Figure 3b), only 0.8% were shared with suburban populations (Figure Sla). In addition, XP-nSL scores appeared less clustered and lower in magnitude among the urban-suburban comparison (Figure Slb) than the suburban-rural (Figure Sla) or urban-rural (Figure 3b) comparisons.

### Text S2: Sweep signatures excluding structured populations

While overall levels of population structure were quite low, one urban and one rural population were outliers along the first two PCs, had slightly elevated pairwise population *F_ST_* when compared to other populations, and clustered separately into their own ancestry categories in the admixture plot (Figure 2). Therefore, we wanted to confirm that our inferred signatures of selective sweeps were not being driven solely by these outlier populations. To do this, we re-ran our selective sweep analysis between urban and rural populations in *selscan* but excluded these populations. 92% of the top urban outlier windows in the original analysis (i.e., those that are outliers relative to the permuted distribution) were also outliers in this revised analysis, while 90% of the top rural outlier windows were shared between these analyses (Figure S2). This suggest that the regions of the genome showing the strongest evidence of selection in urban and rural habitats are not being driven by these population structure outlier populations.

## Supplementary Figures

**Figure S1:**
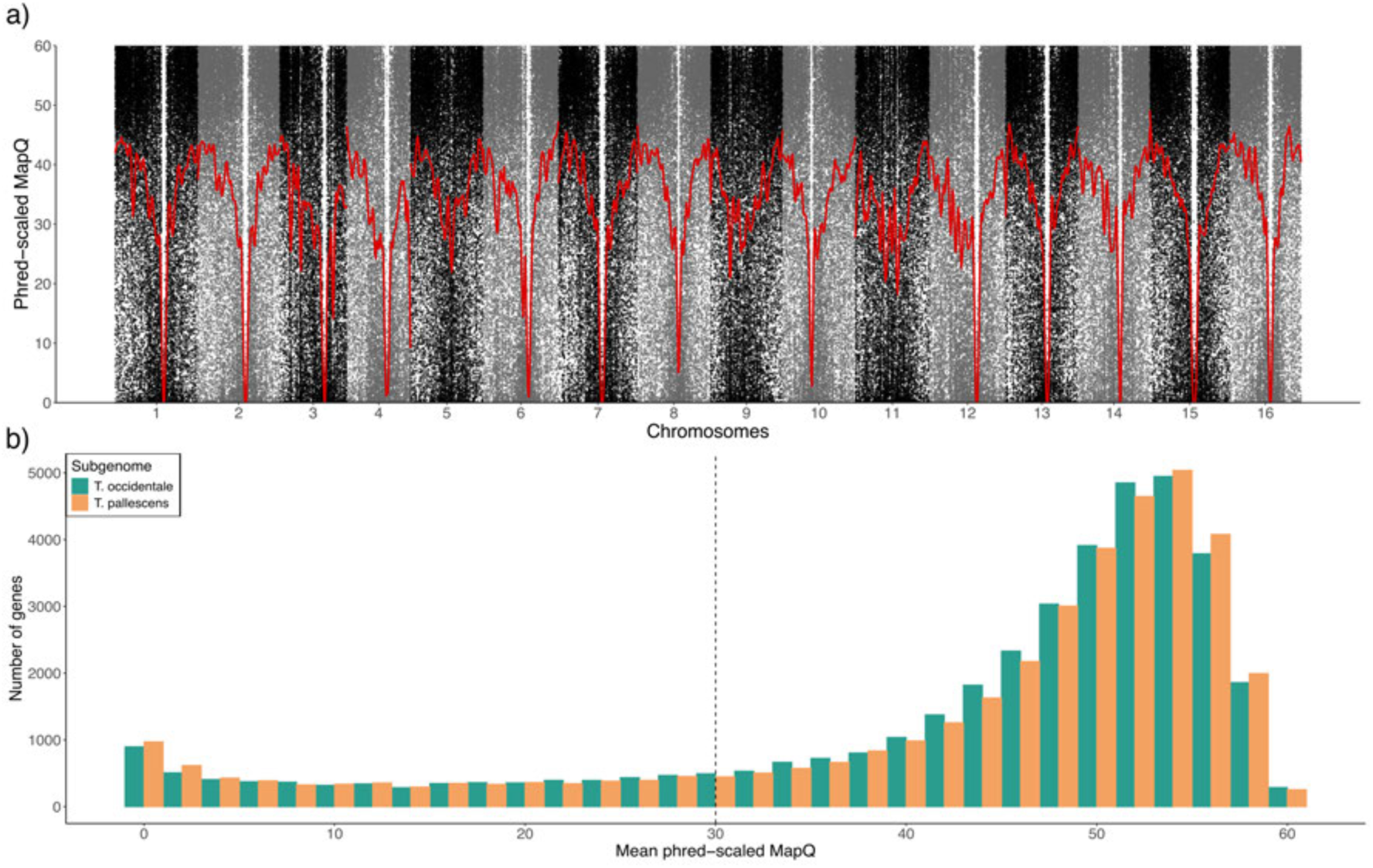
(a) Phred-scaled mapping quality (mapq) scores of reads in 1000 bp windows across the genome with a 200 bp step. Values were LOESS smoothed within chromo­ somes. Odd numbered chromosomes originate from *T. occidentale* while even chromo­ somes originate from *T. pallescens.* (b) Mean mapq scores in genes across both white clover subgenomes. The vertical dashed line represents the phred-scaled mapq threshold applied during read filtering, where reads with mapq < 30 were excluded from all analyses.

**Figure S2:**
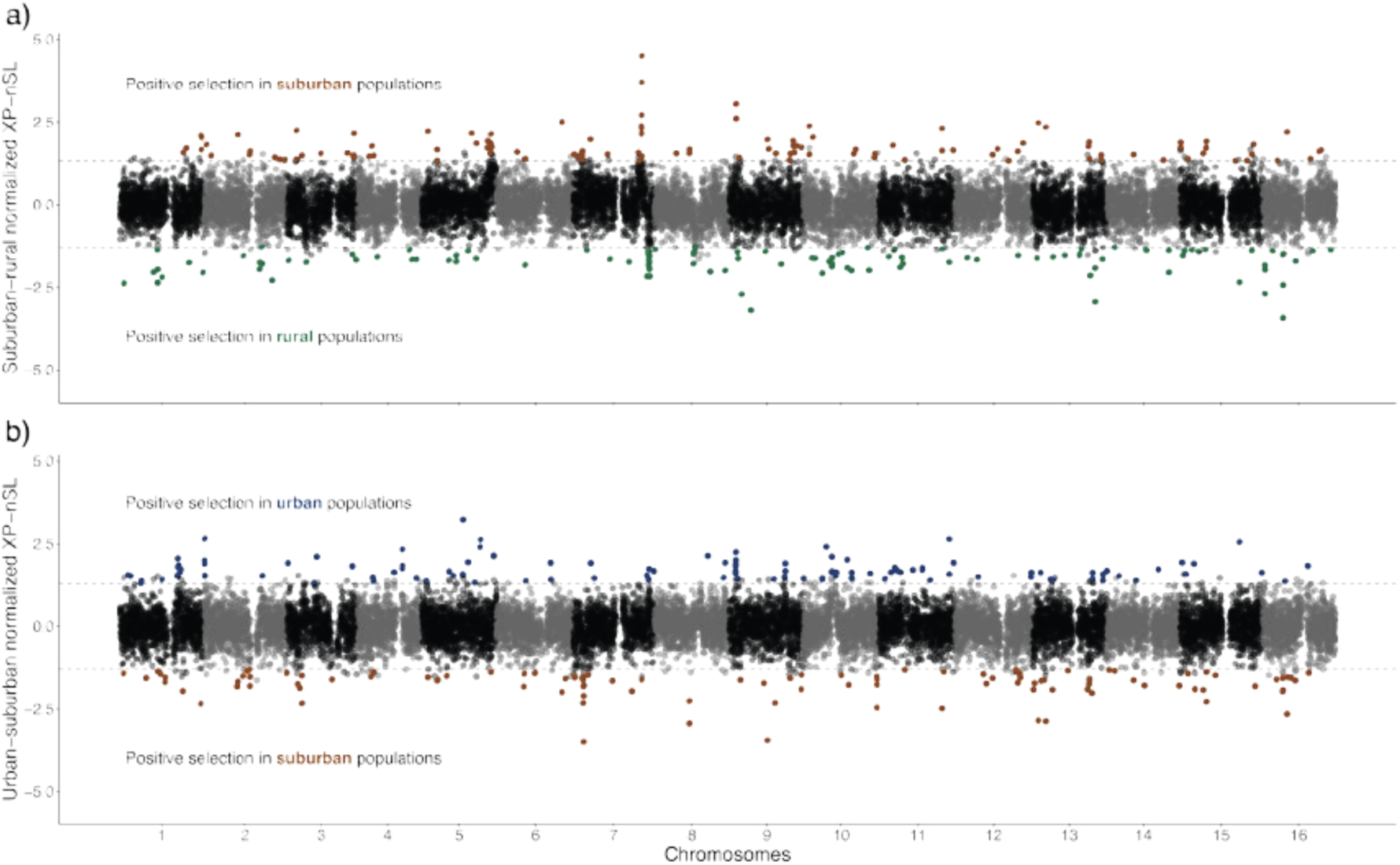
(a) Suburban-rural normalized XP-nSL estimated in 50 kb windows across the genome. Brown points represent outliers showing evidence of positive selection in suburban habitats, while green points represent windows showing evidence of positive selection in rural habitats. (b) Urban-suburban normalized XP-nSL estimated in 50 kb windows across the genome. Blue points represent outliers showing evidence of positive selection in urban habitats, while brown points represent windows showing evidence of positive selection in suburban habitats.

**Figure S3:**
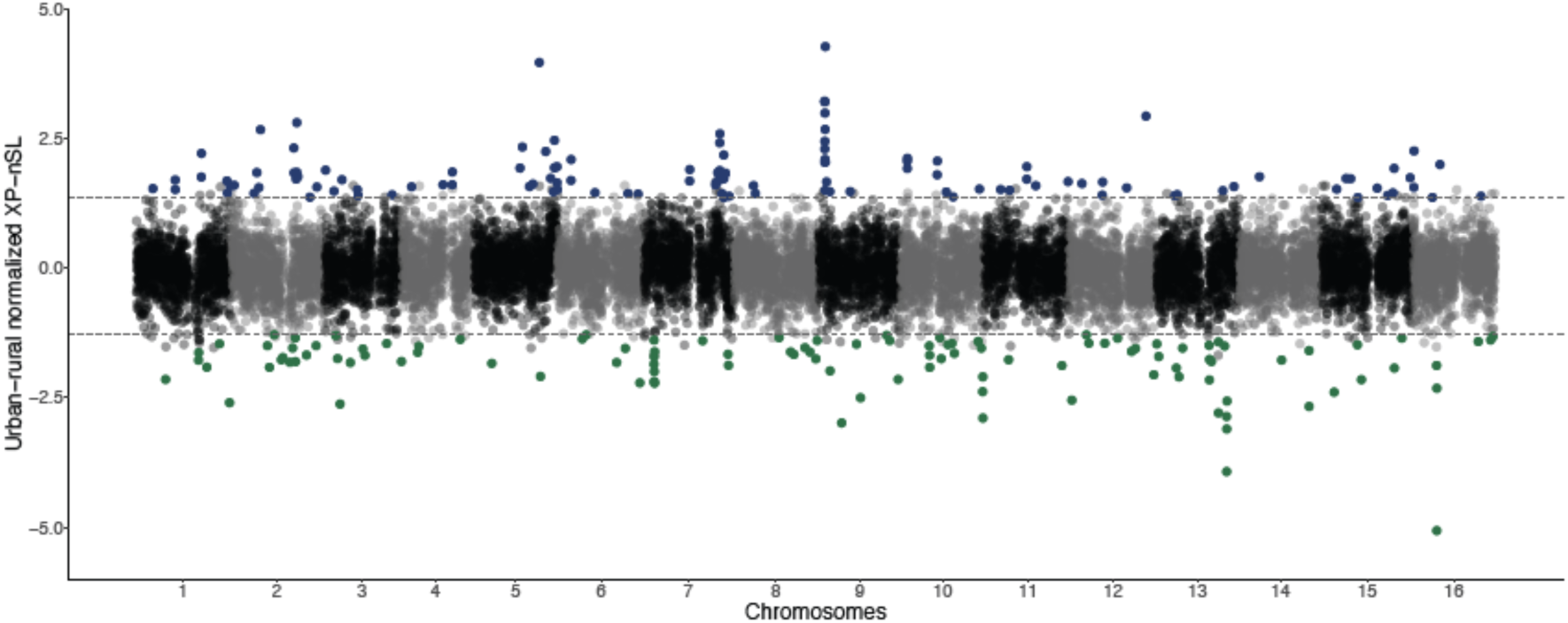
Urban-rural normalized XP-nSL estimated in 50 kb windows across the genome, excluding populations that were outliers in the population structure analyses (Figure 2). Blue points represent outliers showing evidence of positive selection in urban habitats, while green points represent windows showing evidence of positive selection in rural habi­ tats. The Manhattan plot is largely congruent with the one including all populations (Figure 3c), suggest that the strongest signatures of selection are not being driven by these population structure outlier populations.

**Figure S4:**
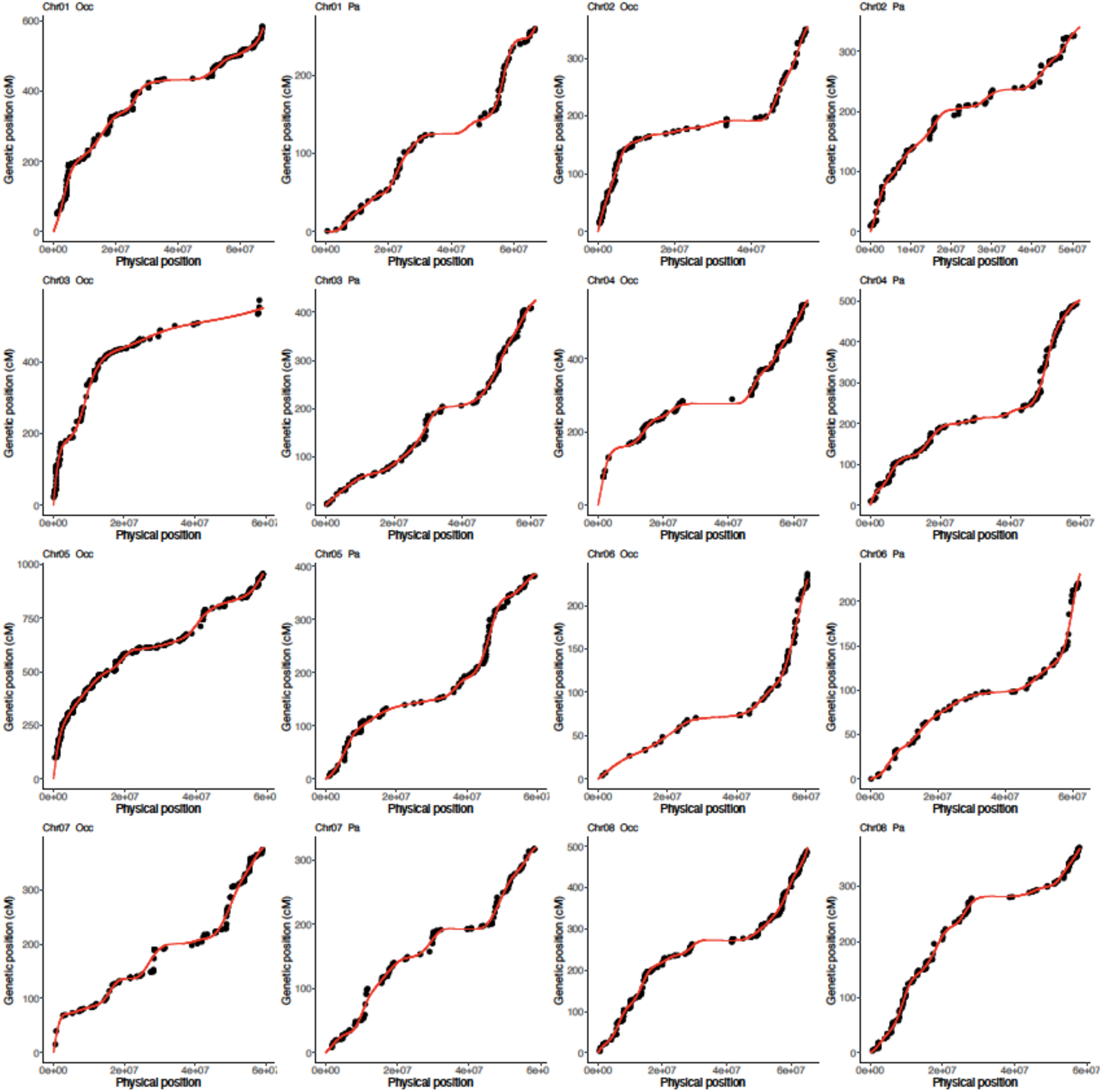
Chromosomal Marey Maps for all 16 white clover chromosomes, fit with inter- polated and monotonically increasing cubic p-splines with 20 knots.

**Figure S5:**
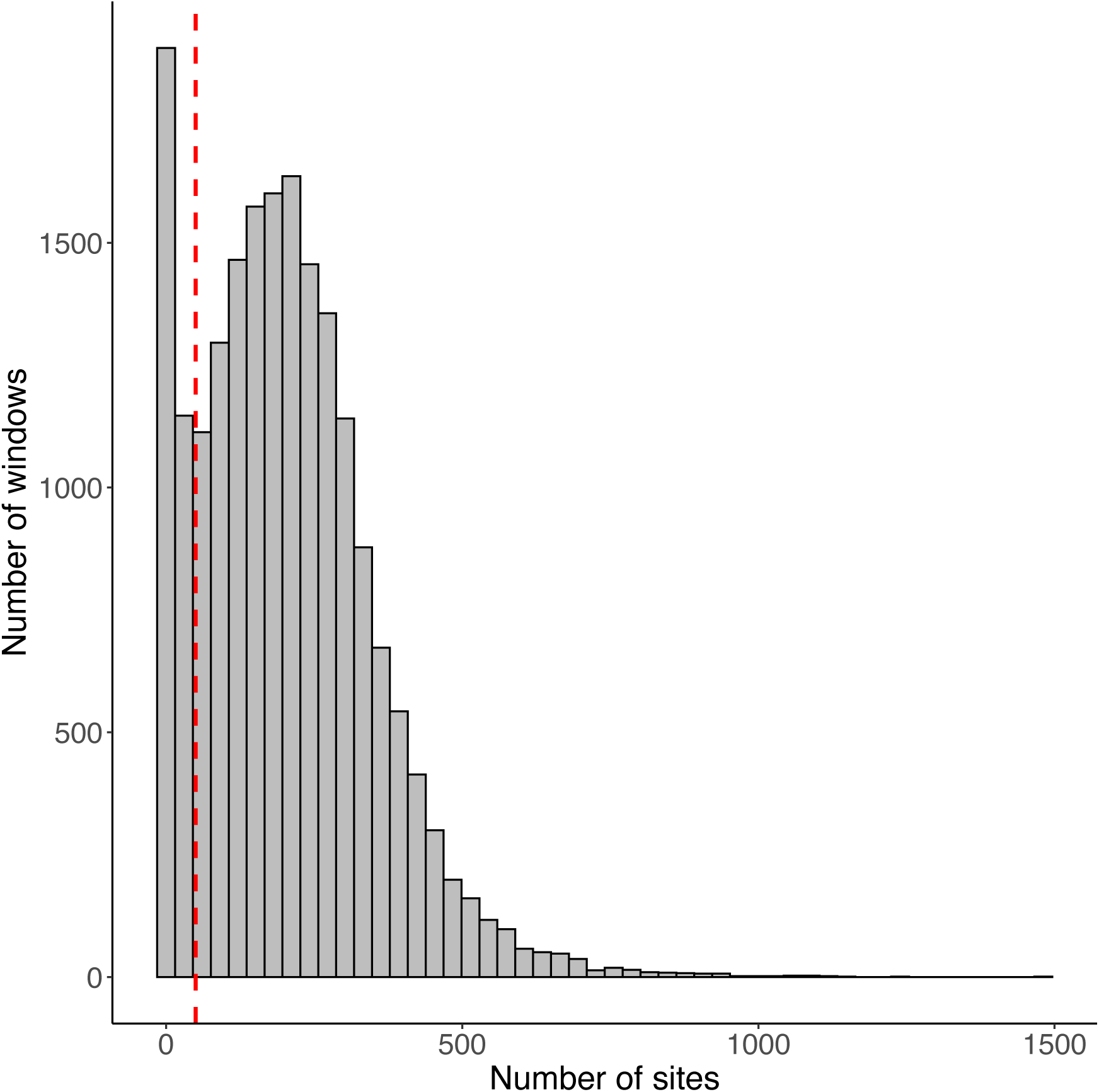
Histogram showing the number of SNPs in all 19,372, 50 kb urban-rural XP­ nSL windows. The vertical red dashed line is at 50 SNPs and represents our cutoff for inclusion of windows. Windows with fewer than 50 SNPs (i.e., left of red line, N = 2,733 or 14.1%) were removed to avoid noisy XP-nSL estimates in regions of low diversity.

**Figure S6:**
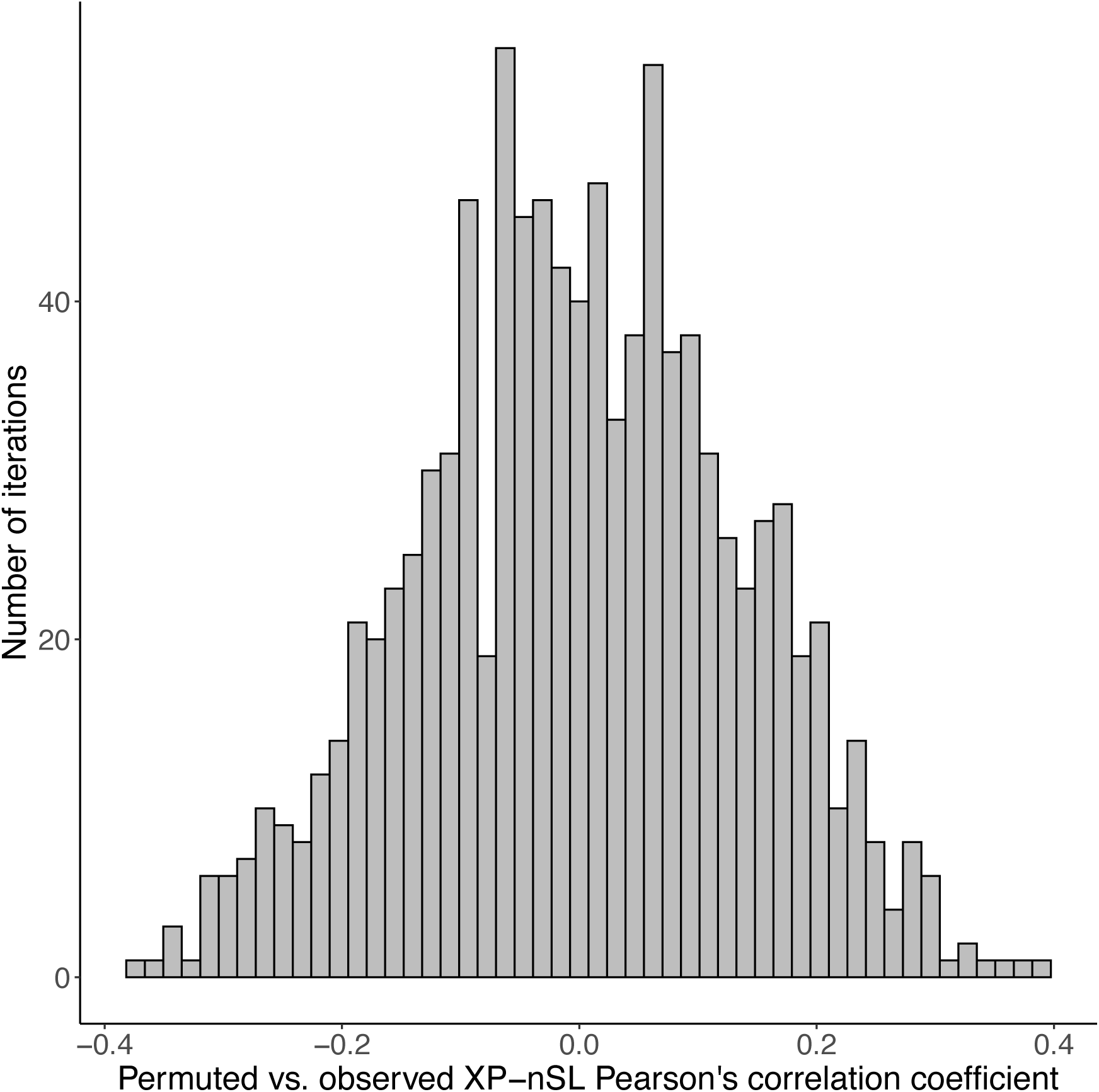
Histogram showing the Pearson’s correlation coefficient comparing observed vs. permuted windowed mean XP-nSL scores across all 1000 permutations of urban and rural samples. As expected, the distribution is centered around zero.

**Figure S7:**
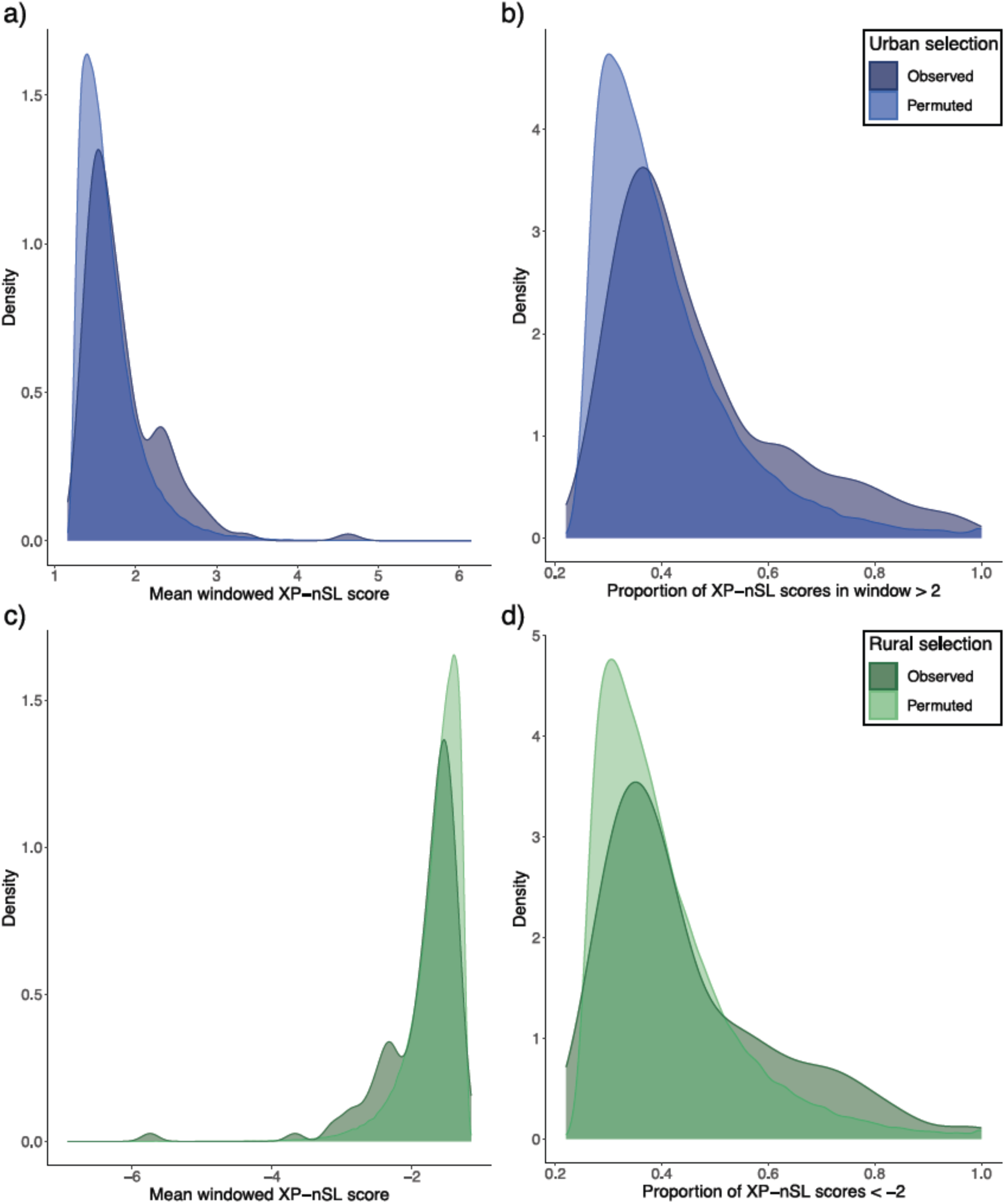
Comparison of observed urban-rural XP-nSL scores to those generated following 1000 permutations of urban and rural samples. (a) Comparison of mean windowed XP-nSL scores for windows with evidence of positive selection in urban habitats. (b) Comparison of the proportion of raw XP-nSL scores with values > 2 for windows with evidence of positive selection in urban habitats. Panels (c) and (d) are the same as (a) and (b), but for windows showing evidence of positive selection in rural habitats.

**Figure S8:**
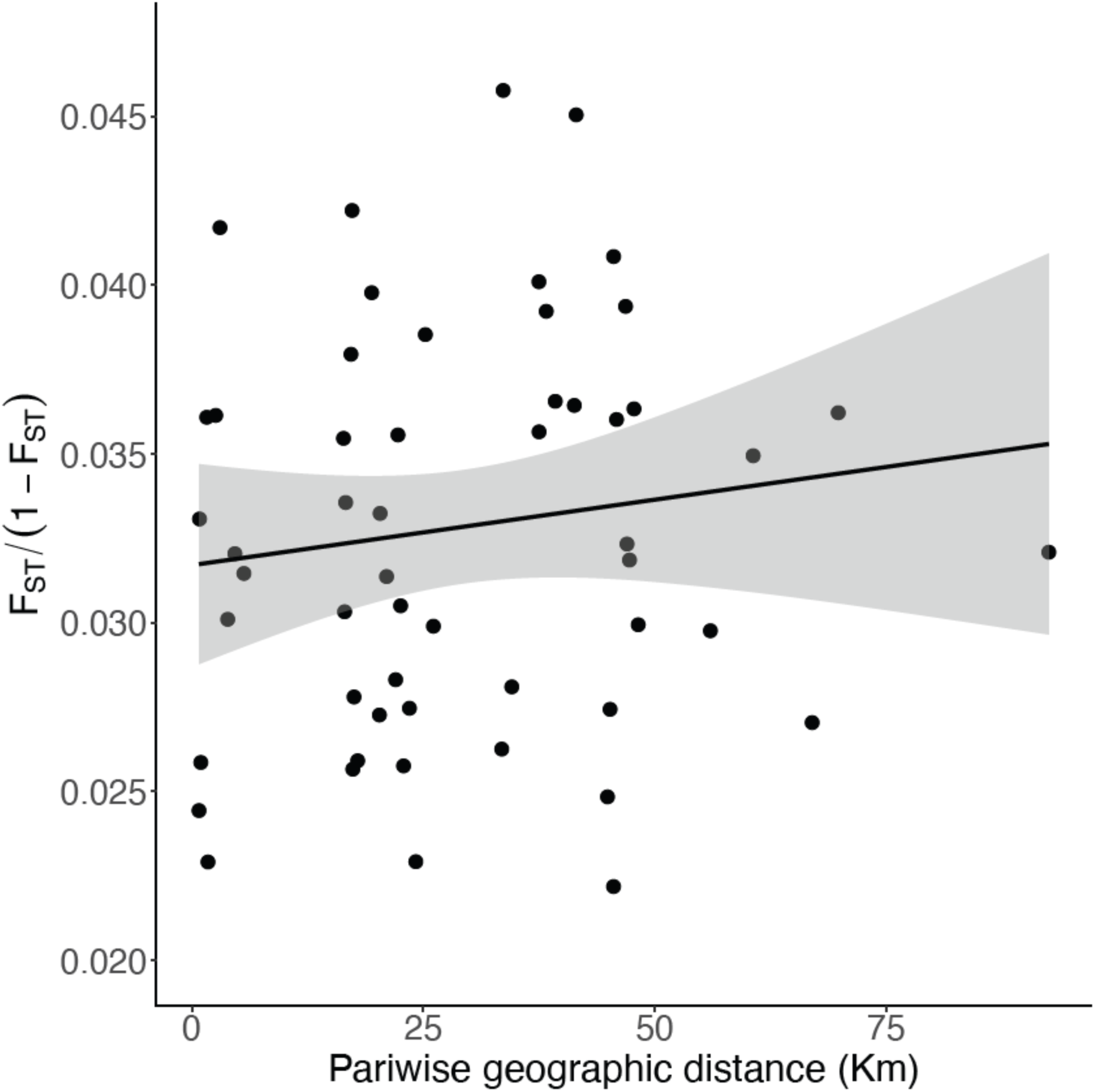
Scatterplot showing the relationship between linearized *F_ST_* and geographic distance between all populations containing more than one individual (all N >= 7). We did not detect significant evidence of isolation by distance at this scale (Mantel r = 0.13, *P* = 0.26).

**Figure S9:**
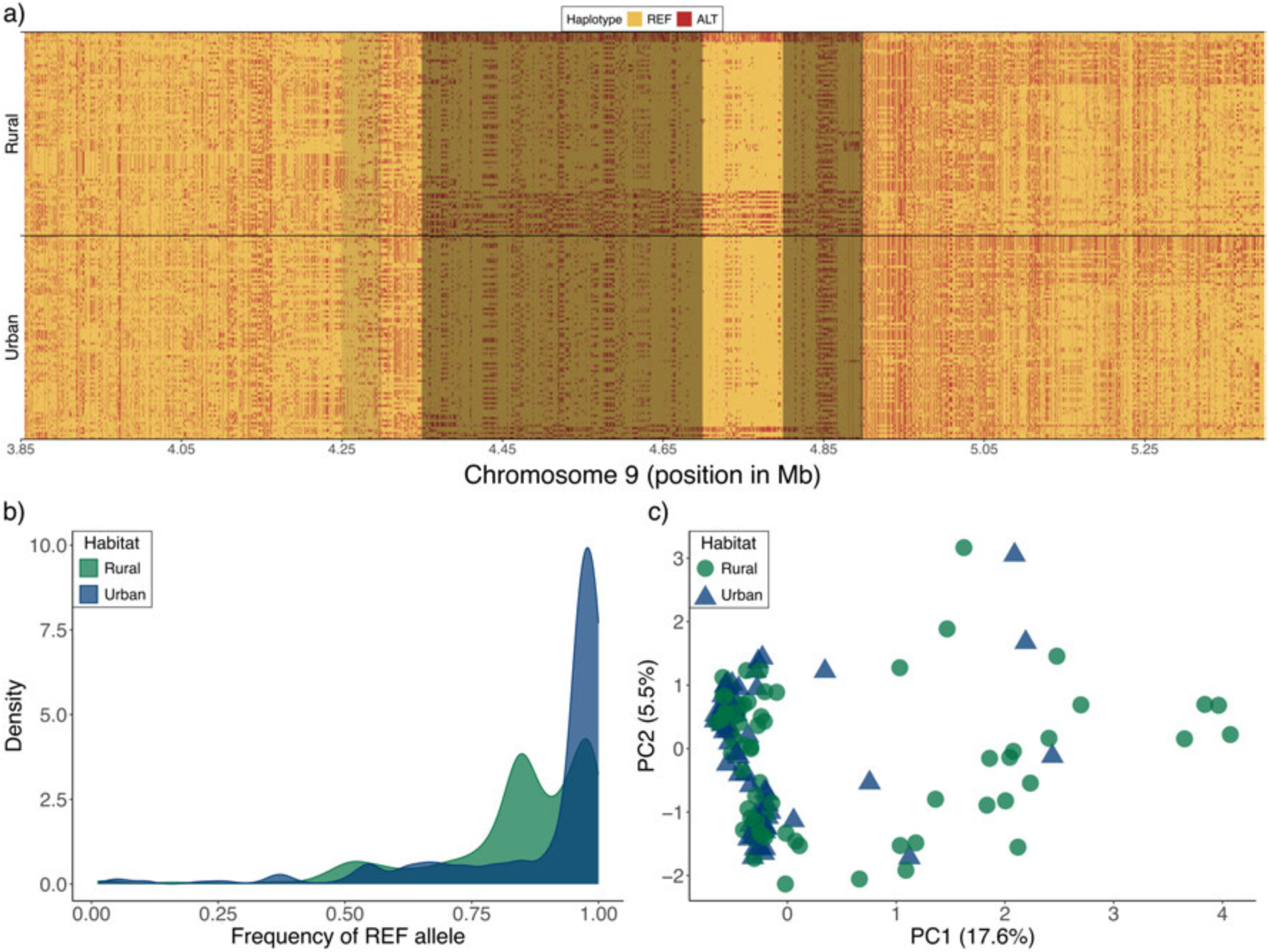
(a) Raw haplotypes in urban and rural habitats for region shown in Figure 4c with evidence of positive selection in rural habitats. Each individual is represented by two rows (one for each haplotype), with reference alleles coloured yellow and alternative alleles colored red. Light grey bands display XP-nSL outlier windows, while dark grey bands are XP-nSL windows that are additionally in the top 5% of the urban-rural permuted XP-nSL distribution. Figure generated using *genotype_plot.R* (Whiting 2022). (b) Frequency of the REF allele in urban and rural populations across all sites highlighted in grey from (a). (c) Principal components analysis performed on haplotype distances for regions highlighted in grey in (a).

**Figure S10:**
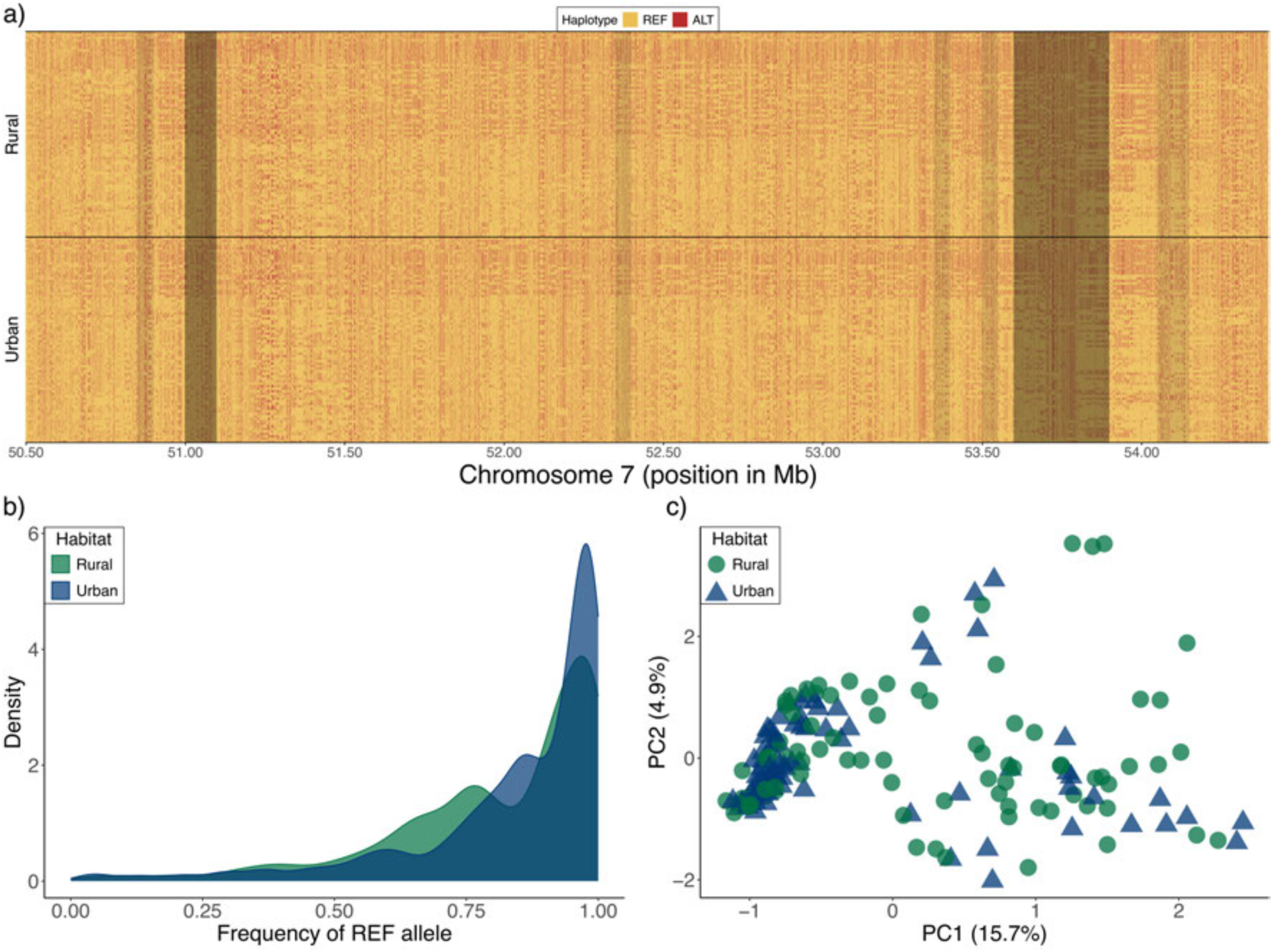
(a) Raw haplotypes in urban and rural habitats for region shown in Figure 4b with evidence of positive selection in urban habitats. Each individual is represented by two rows (one for each haplotype), with reference alleles coloured yellow and alternative alleles colored red. Light grey bands display XP-nSL outlier windows, while dark grey bands are XP-nSL windows that are additionally in the top 5% of the urban-rural permuted XP-nSL distribution. Figure generated using *genotype_plot.R* (Whiting 2022). (b) Frequency of the REF allele in urban and rural populations across all sites highlighted in grey from (a). (c) Principal components analysis performed on haplotype distances for regions highlighted in grey in (a).

**Figure S11:**
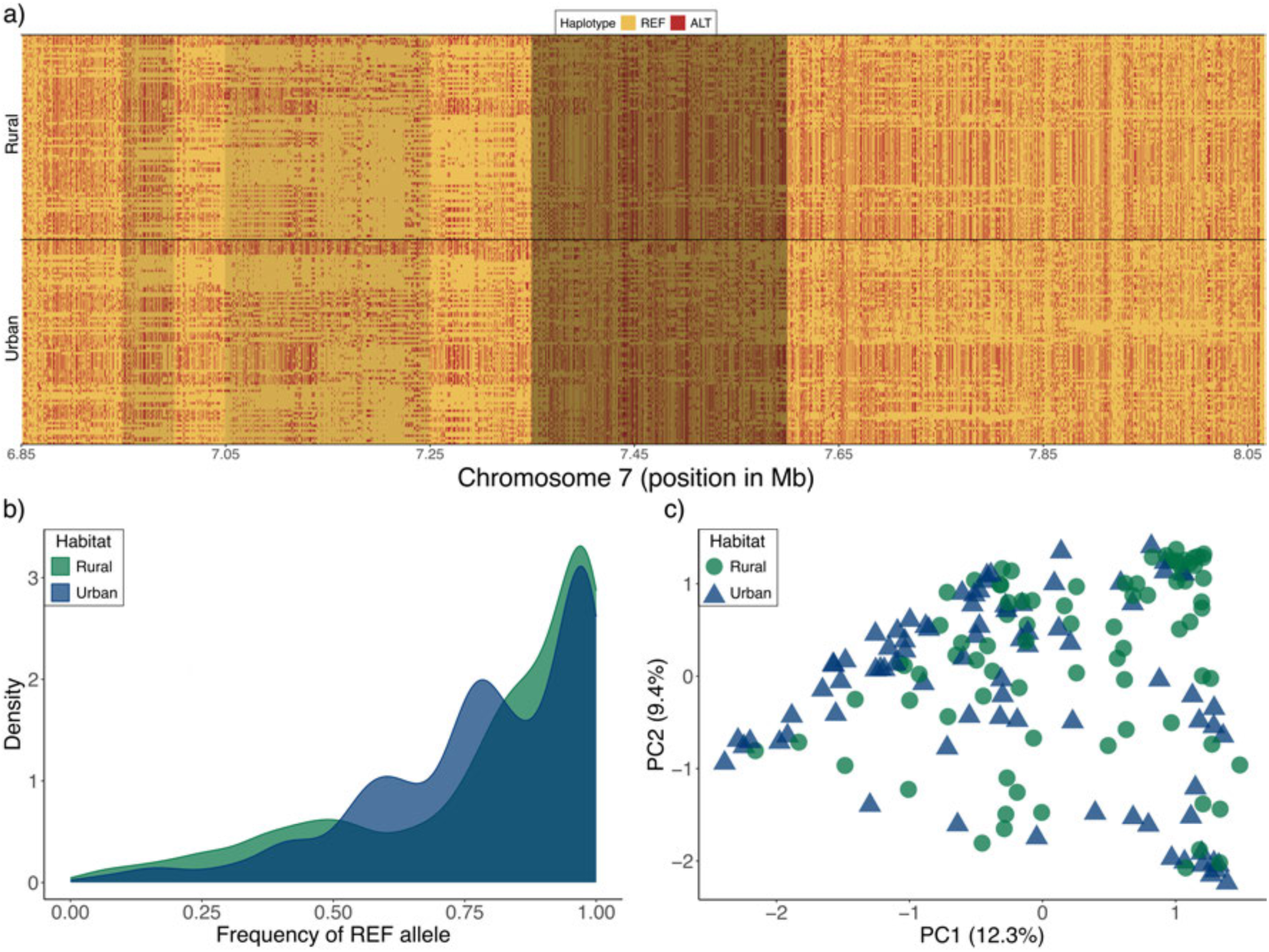
(a) Raw haplotypes in urban and rural habitats for region shown in Figure 4a with evidence of positive selection in urban habitats. Each individual is represented by two rows (one for each haplotype), with reference alleles coloured yellow and alternative alleles colored red. Light grey bands display XP-nSL outlier windows, while dark grey bands are XP-nSL windows that are additionally in the top 5% of the urban-rural permuted XP-nSL distribution. Figure generated using *genotype_plot.R* (Whiting 2022). (b) Frequency of the REF allele in urban and rural populations across all sites highlighted in grey from (a). (c) Principal components analysis performed on haplotype distances for regions highlighted in grey in (a).

**Figure S12:**
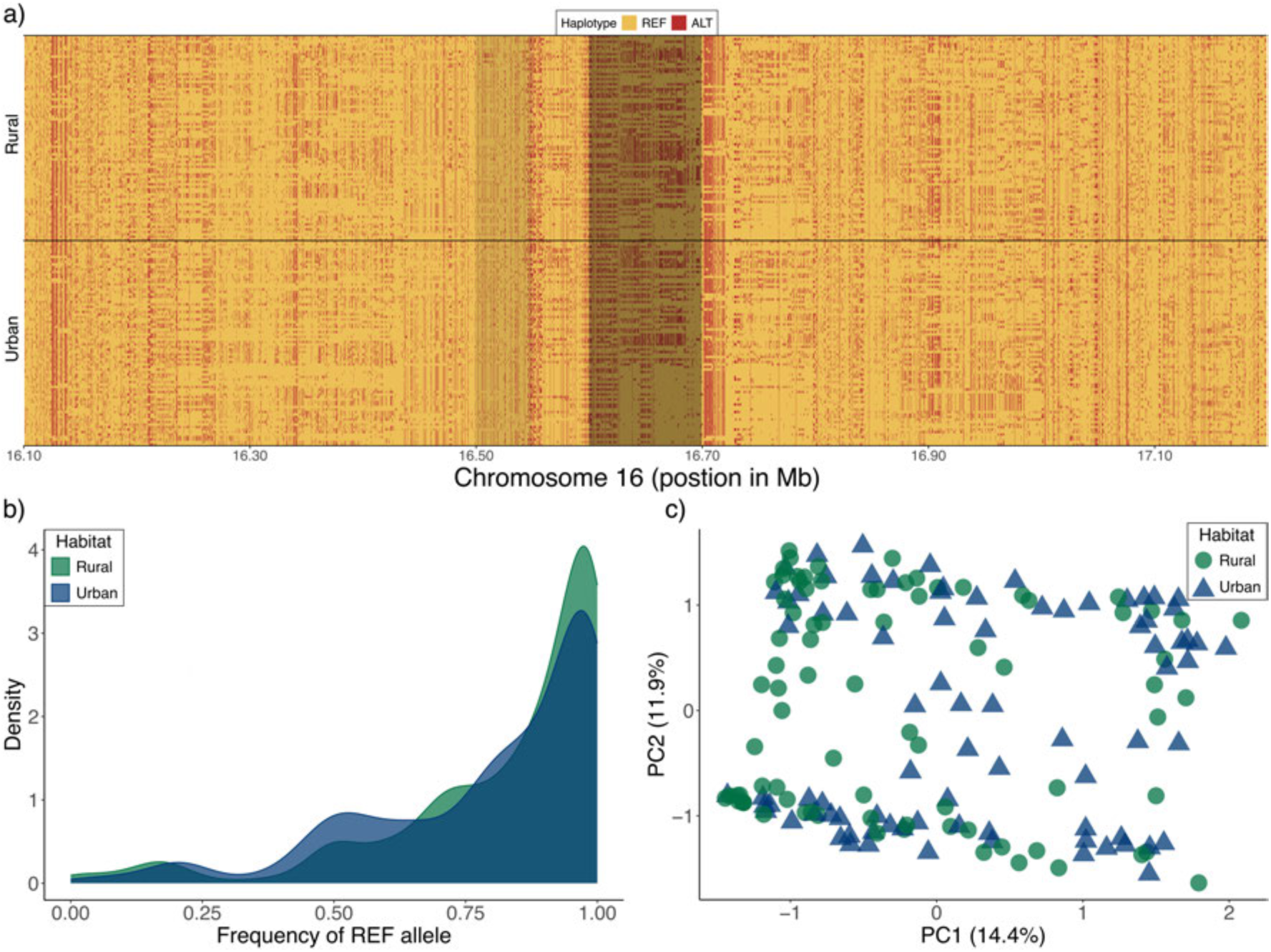
(a) Raw haplotypes in urban and rural habitats for region shown in Figure 4d with evidence of positive selection in urban habitats. Each individual is represented by two rows (one for each haplotype), with reference alleles coloured yellow and alternative alleles colored red. Light grey bands display XP-nSL outlier windows, while dark grey bands are XP-nSL windows that are additionally in the top 5% of the urban-rural permuted XP-nSL distribution. Figure generated using *genotype_plot.R* (Whiting 2022). (b) Frequency of the REF allele in urban and rural populations across all sites highlighted in grey from (a). (c) Principal components analysis performed on haplotype distances for regions highlighted in grey in (a).

**Figure S13:**
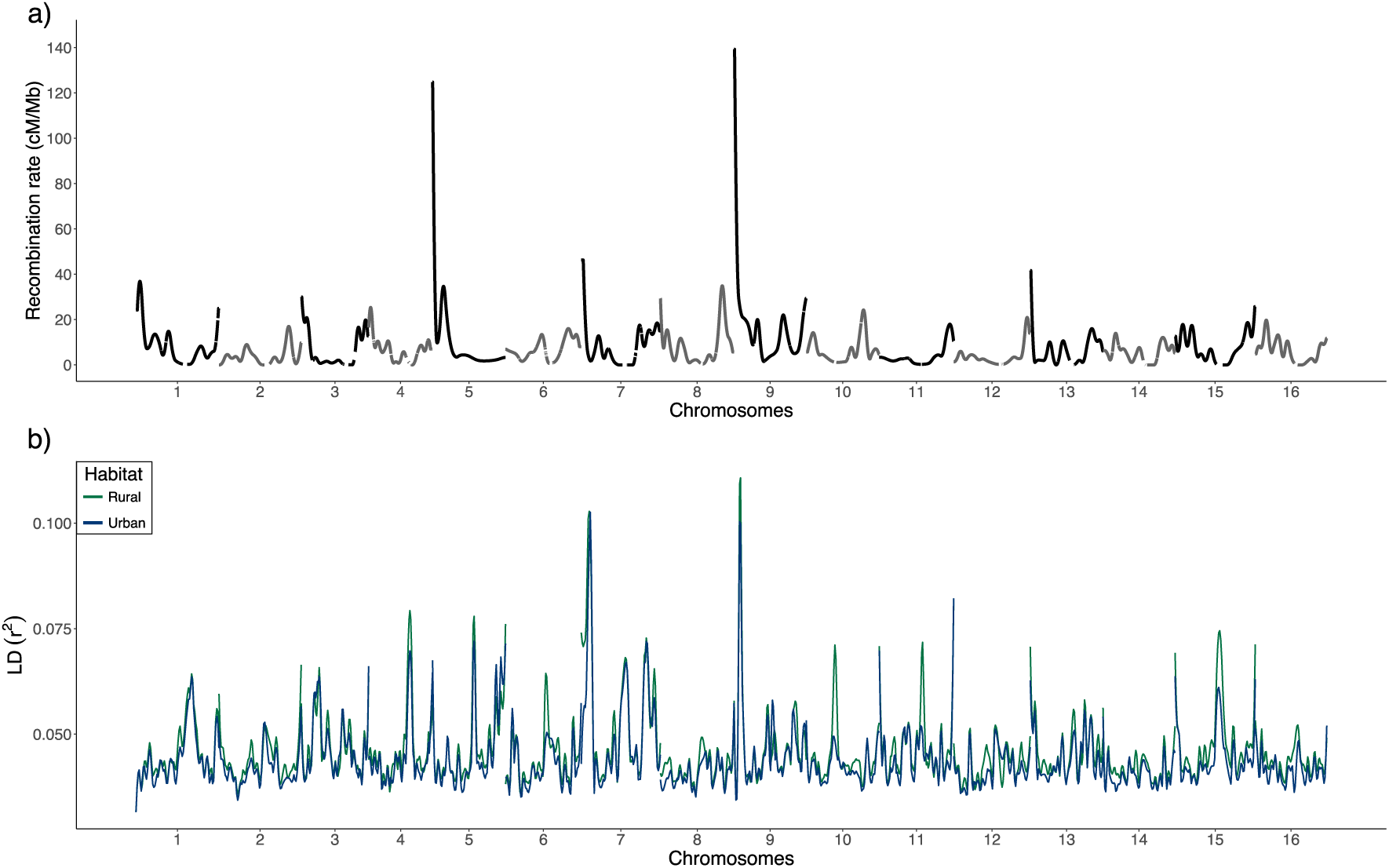
(a) Genome-wide recombination (in cM/mB) generated using the interpolated genetic map from Figure S3. The map was LOESS-smoothed within chromosomes. (b) Windowed LD (r^2^) in urban (blue) and rural (green) populations. LD was estimated as the mean *r^2^* in 50 kb windows using PLINK v.1.9.

**Figure S14:**
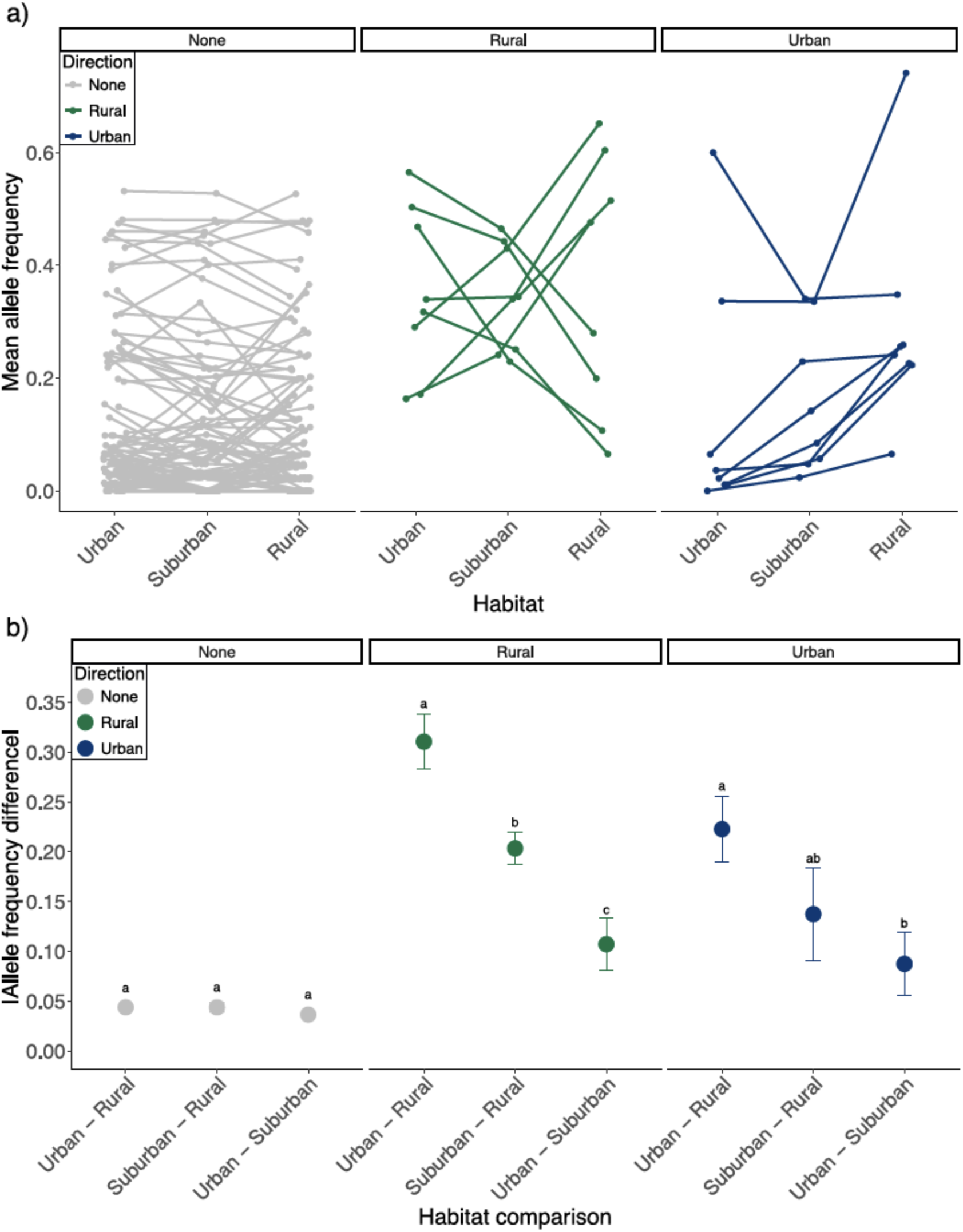
(a) Minor allele frequency at the most differentiated SNP in each of the top candidate regions of selection (Table 1) and in each habitat. Lines are colored based on the inferred direction of selection from the XP-nSL analysis with blue lines indicating positive selection in urban habitats and green lines reflecting positive selection in rural habitats. Grey lines show allele frequencies for 100 randomly selected SNPs across the genome. (b) Mean absolute value of the urban-rural, surburban-rural, and urban-suburban difference in allele frequencies across the most differentiated SNPs from the top selected regions (Table 1). Different letters within panels denote significant differences at *P* < 0.05 from Tu.key HSD tests.

**Figure S15:**
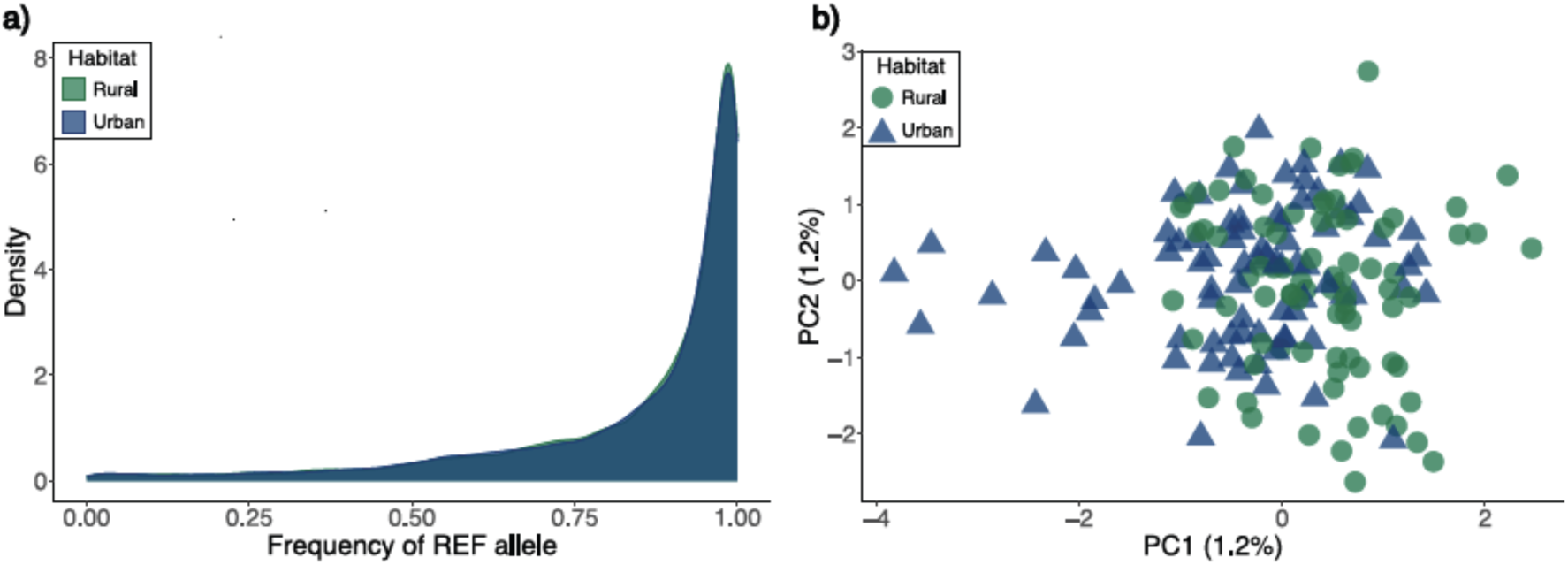
Urban and rural allele frequencies and haplotype differentiation across 100 randomly-selected, unselected regions (i.e., not XP-nSL outliers) along chromosome 1. (a) Frequency of the REF allele in urban and rural populations across all sites. Note that urban and rural allele frequency histograms are perfectly overlapping. (b) Principal components analysis performed on haplotype distances across regions.

**Figure S16:**
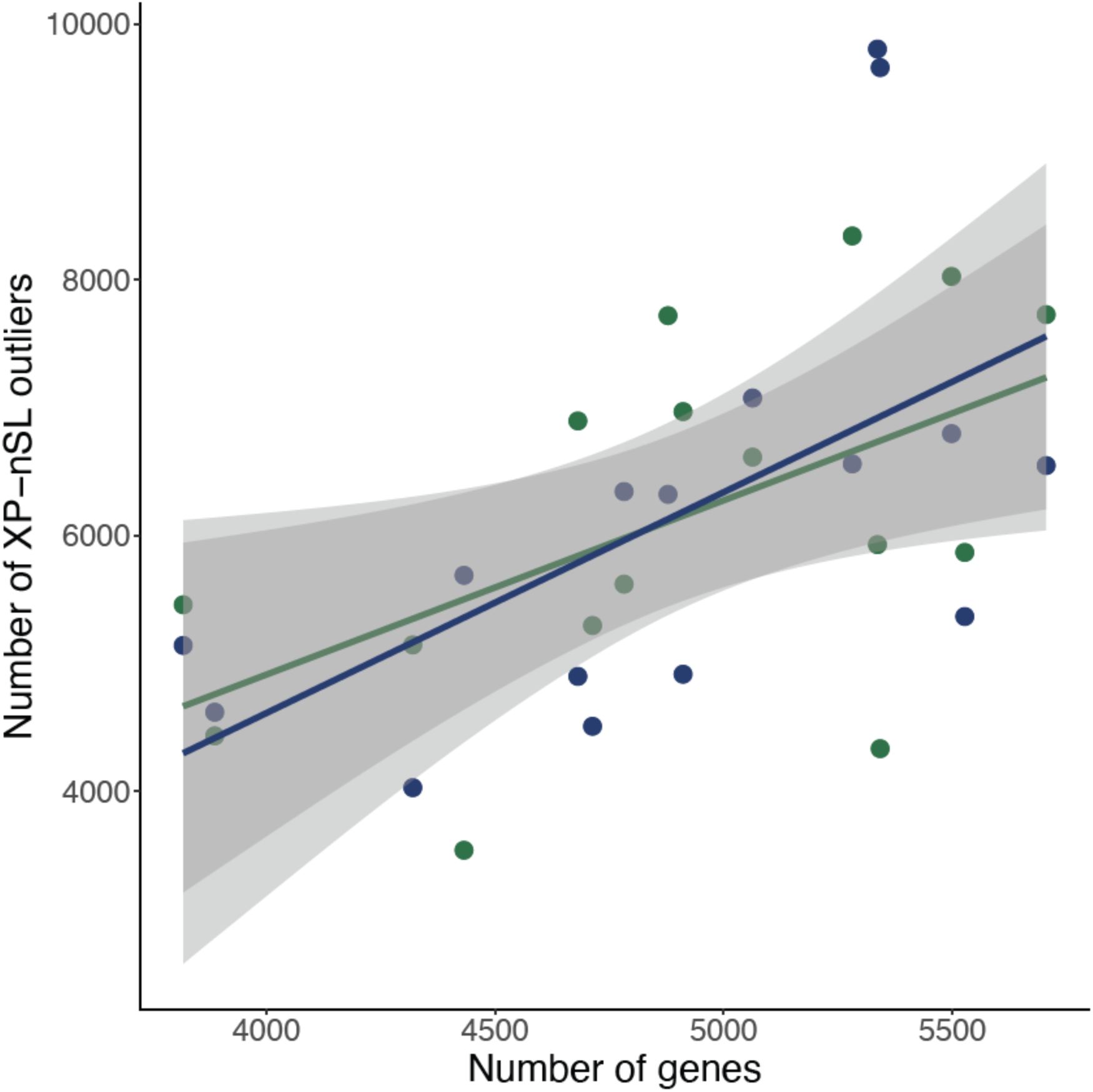
Relationship between the number of genes on a chromosome and the number of XP-nSL outlier loci in both urban (blue) and rural (green) habitats. The number of genes is a significant predictor of the number of outliers *(P* = 0.03), with no significant difference in this effect between habitats (# of genes *x* # of outliers interaction, *P* = 0.67

## Supplementary Tables

**Table S1:** Significant Gene Ontology (GO) categories for Biological Processes for genes in candidate regions of selective sweeps in urban and rural populations.

| GO.ID | Term | Annotated | Significant | Expected | Pval | Habitat |
| --- | --- | --- | --- | --- | --- | --- |
| GO:0015693 | magnesium ion transport | 22 | 2 | 0.04 | 0.00059 | Urban |
| GO:0000972 | transcription-dependent tethering of RNA... | 29 | 2 | 0.05 | 0.00103 | Urban |
| GO:0051083 | 'de novo' cotranslational protein foldin... | 7 | 1 | 0.01 | 0.01137 | Urban |
| GO:0006546 | glycine catabolic process | 8 | 1 | 0.01 | 0.01298 | Urban |
| GO:0006405 | RNA export from nucleus | 108 | 2 | 0.18 | 0.01351 | Urban |
| GO:0006450 | regulation of translational fidelity | 9 | 1 | 0.01 | 0.01459 | Urban |
| GO:0006096 | glycolytic process | 116 | 2 | 0.19 | 0.01547 | Urban |
| GO:0034067 | protein localization to Golgi apparatus | 10 | 1 | 0.02 | 0.0162 | Urban |
| GO:0010089 | xylem development | 11 | 1 | 0.02 | 0.0178 | Urban |
| GO:0006606 | protein import into nucleus | 127 | 2 | 0.21 | 0.01834 | Urban |
| GO:0006357 | regulation of transcription by RNA polym... | 956 | 5 | 1.56 | 0.02065 | Urban |
| GO:0019264 | glycine biosynthetic process from serine | 16 | 1 | 0.03 | 0.02579 | Urban |
| GO:0071586 | CAAX-box protein processing | 16 | 1 | 0.03 | 0.02579 | Urban |
| GO:0035999 | tetrahydrofolate interconversion | 16 | 1 | 0.03 | 0.02579 | Urban |
| GO:0006047 | UDP-N-acetylglucosamine metabolic proces... | 19 | 1 | 0.03 | 0.03056 | Urban |
| GO:0006623 | protein targeting to vacuole | 29 | 1 | 0.05 | 0.04627 | Urban |
| GO:0007021 | tubulin complex assembly | 7 | 2 | 0.01 | 5e-05 | Rural |
| GO:0007023 | post-chaperonin tubulin folding pathway | 10 | 2 | 0.02 | 0.00011 | Rural |
| GO:0045492 | xylan biosynthetic process | 42 | 2 | 0.07 | 0.00199 | Rural |
| GO:0009057 | macromolecule catabolic process | 1618 | 3 | 2.54 | 0.00675 | Rural |
| GO:0032784 | regulation of DNA-templated transcriptio... | 91 | 2 | 0.14 | 0.00904 | Rural |
| GO:0016559 | peroxisome fission | 6 | 1 | 0.01 | 0.00938 | Rural |
| GO:0032957 | inositol trisphosphate metabolic process | 11 | 1 | 0.02 | 0.01712 | Rural |
| GO:0046168 | glycerol-3-phosphate catabolic process | 12 | 1 | 0.02 | 0.01867 | Rural |
| GO:0030490 | maturation of SSU-rRNA | 138 | 2 | 0.22 | 0.01991 | Rural |
| GO:0007018 | microtubule-based movement | 140 | 2 | 0.22 | 0.02045 | Rural |
| GO:0006376 | mRNA splice site selection | 15 | 1 | 0.02 | 0.02328 | Rural |
| GO:0007059 | chromosome segregation | 249 | 2 | 0.39 | 0.03502 | Rural |
| GO:0045039 | protein insertion into mitochondrial inn... | 24 | 1 | 0.04 | 0.03699 | Rural |
| GO:0006282 | regulation of DNA repair | 27 | 1 | 0.04 | 0.04152 | Rural |

**Table S2:** Significant Gene Ontology (GO) categories for Molecular Function for genes in candidate regions of selective sweeps in urban and rural populations.

| GO.ID | Term | Annotated | Significant | Expected | Pval | Habitat |
| --- | --- | --- | --- | --- | --- | --- |
| GO:0015095 | magnesium ion transmembrane transporter ... | 22 | 2 | 0.03 | 0.00054 | Urban |
| GO:0000978 | RNA polymerase II cis-regulatory region ... | 439 | 5 | 0.68 | 0.00061 | Urban |
| GO:0030544 | Hsp70 protein binding | 25 | 2 | 0.04 | 0.0007 | Urban |
| GO:0030955 | potassium ion binding | 28 | 2 | 0.04 | 0.00088 | Urban |
| GO:0004743 | pyruvate kinase activity | 28 | 2 | 0.04 | 0.00088 | Urban |
| GO:0017056 | structural constituent of nuclear pore | 47 | 2 | 0.07 | 0.00246 | Urban |
| GO:0046983 | protein dimerization activity | 982 | 6 | 1.53 | 0.0042 | Urban |
| GO:0033612 | receptor serine/threonine kinase binding | 70 | 2 | 0.11 | 0.00537 | Urban |
| GO:0020037 | heme binding | 1103 | 6 | 1.72 | 0.0073 | Urban |
| GO:0045296 | cadherin binding | 5 | 1 | 0.01 | 0.00775 | Urban |
| GO:0004412 | homoserine dehydrogenase activity | 9 | 1 | 0.01 | 0.01391 | Urban |
| GO:0005506 | iron ion binding | 955 | 5 | 1.49 | 0.01644 | Urban |
| GO:0004072 | aspartate kinase activity | 13 | 1 | 0.02 | 0.02003 | Urban |
| GO:0016866 | intramolecular transferase activity | 176 | 3 | 0.27 | 0.02094 | Urban |
| GO:0004372 | glycine hydroxymethyltransferase activit... | 16 | 1 | 0.02 | 0.0246 | Urban |
| GO:0004497 | monooxygenase activity | 902 | 5 | 1.4 | 0.02861 | Urban |
| GO:0016705 | oxidoreductase activity, acting on paire... | 940 | 5 | 1.46 | 0.0334 | Urban |
| GO:0016787 | hydrolase activity | 7781 | 15 | 12.1 | 0.03787 | Urban |
| GO:0016836 | hydro-lyase activity | 198 | 2 | 0.31 | 0.0382 | Urban |
| GO:0016868 | intramolecular transferase activity, pho... | 31 | 1 | 0.05 | 0.04712 | Urban |
| GO:0008194 | UDP-glycosyltransferase activity | 525 | 3 | 0.82 | 0.04853 | Urban |
| GO:0048487 | beta-tubulin binding | 5 | 2 | 0.01 | 1.9e-05 | Rural |
| GO:0015018 | galactosylgalactosylxylosylprotein 3-bet... | 10 | 2 | 0.01 | 8.7e-05 | Rural |
| GO:0042285 | xylosyltransferase activity | 14 | 2 | 0.02 | 0.00018 | Rural |
| GO:0005524 | ATP binding | 4687 | 16 | 6.59 | 0.00055 | Rural |
| GO:0016887 | ATP hydrolysis activity | 1201 | 7 | 1.69 | 0.00144 | Rural |
| GO:0010333 | terpene synthase activity | 68 | 2 | 0.1 | 0.00417 | Rural |
| GO:0000287 | magnesium ion binding | 265 | 3 | 0.37 | 0.00624 | Rural |
| GO:0003724 | RNA helicase activity | 125 | 2 | 0.18 | 0.01349 | Rural |
| GO:0008017 | microtubule binding | 365 | 3 | 0.51 | 0.01482 | Rural |
| GO:0047325 | inositol tetrakisphosphate 1-kinase acti... | 11 | 1 | 0.02 | 0.01537 | Rural |
| GO:0052725 | inositol-1,3,4-trisphosphate 6-kinase ac... | 11 | 1 | 0.02 | 0.01537 | Rural |
| GO:0052726 | inositol-1,3,4-trisphosphate 5-kinase ac... | 11 | 1 | 0.02 | 0.01537 | Rural |
| GO:0003777 | microtubule motor activity | 140 | 2 | 0.2 | 0.01671 | Rural |
| GO:0047952 | glycerol-3-phosphate dehydrogenase [NAD(... | 12 | 1 | 0.02 | 0.01676 | Rural |
| GO:0003917 | DNA topoisomerase type I (single strand ... | 16 | 1 | 0.02 | 0.02228 | Rural |
| GO:0019139 | cytokinin dehydrogenase activity | 19 | 1 | 0.03 | 0.0264 | Rural |
| GO:0140323 | solute:anion antiporter activity | 21 | 1 | 0.03 | 0.02914 | Rural |
| GO:0003688 | DNA replication origin binding | 22 | 1 | 0.03 | 0.03051 | Rural |
| GO:0005388 | P-type calcium transporter activity | 28 | 1 | 0.04 | 0.03867 | Rural |
| GO:0004332 | fructose-bisphosphate aldolase activity | 35 | 1 | 0.05 | 0.0481 | Rural |
| GO:0015605 | organophosphate ester transmembrane tran... | 36 | 1 | 0.05 | 0.04944 | Rural |

## References

Alexa, A. and J. Rahnenfuhrer. 2016. topGO: Enrichment Analysis for Gene On­ tology. Version 2.50.0.

Andrews, S. et al. 2010. FastQC: a quality control tool for high throughput sequence data.

Baker, H. G. 1974. The evolution of weeds. Annual Review of Ecology_1_ Evolution_1_ and Systematics 5: 1–24.

Barnett, D. W., E. K. Garrison, A. R. Quinlan, M. P. Stromberg, and G. T. Marth. 2011. BamTools: a C++ API and toolkit for analyzing and managing BAM files. Bioinformatics 27: 1691–1692.

Bartlewicz, J., K. Vandepitte, H. Jacquemyn, and O. Hannay. 2015. Population genetic diversity of the clonal self-incompatible herbaceous plant *Linaria vulgaris* along an urbanization gradient. Biol. J. Linn. Soc. Land. 116: 603–613.

Battlay, P., B. T. Hendrickson, J. I. Mendez-Reneau, J. S. Santangelo, L. J. Albano, J. Wilson, A. E. Caizergues, N. King, A. Puentes, A. Tudoran, C. Violle, F. Vasseur, C. M. Patterson, M. J. Foster, C. Stamps, S. G. Innes, R. Allio, F. Angeoletto, D. N. Anstett, J. Anstett, A. Bucharova, M. S. Comerford, S. David, M. Falahati Anbaran, W. Godsoe, C. Gonzalez-Lagos, P. E. Gundel, G. R. Hood, R. Karousou, C. Lampei, C. Lara, A. Lazaro-Lobo, D. Leandro, T. J. S. Merritt, N. Mitchell, M. Mohammadi Bazargani, A. Moles, M. Murua, J. Paule, V. Pfeiffer, J. A. M. Raeymaekers, D. J. Rennison, R. S. Rios, J. K. Rowntree, A. C. Schneider, K. Stack Whitney, I. Tamburrino, A. VanWallendael, P. Y. Kim, R. W. Ness, M. T. J. Johnson, K. A. Hodgins, and N. J. Kooyers. 2024. Structural variants underlie parallel adaptation following global invasion. Evolutionary Biology.

Beninde, J., S. Feldmeier, M. Veith, and A. Hochkirch. 2018. Admixture of hybrid swarms of native and introduced lizards in cities is determined by the cityscape structure and invasion history. Proceedings of the Royal Society B: Biological Sciences 285: 20180143.

Beninde, J., S. Feldmeier, M. Werner, D. Peroverde, U. Schulte, A. Hochkirch, and M. Veith. 2016. Cityscape genetics: structural vs. functional connectivity of an urban lizard population. Mal. Ecol. 25: 4984–5000.

Bhatia, G., N. Patterson, S. Sankararaman, and A. L. Price. 2013. Estimating and interpreting *F_ST_*: the impact of rare variants. Genome Res. 23: 1514–1521.

Bjorklund, M., I. Ruiz, and J. C. Senar. 2010. Genetic differentiation in the urban habitat: the great tits *(Parus major)* of the parks of Barcelona city. Biol. J. Linn. Soc. Land. 99: 9–19.

Brans, K., R. Stoks, and L. De Meester. 2018. Urbanization drives genetic differ­ entiation in physiology and structures the evolution of pace-of-life syndromes in the water flea *Daphnia magna*. Proceedings of the Royal Society B: Biological Sciences 285: 20180169.

Brans, K. I. and L. De Meester. 2018. City life on fast lanes: Urbanization induces an evolutionary shift towards a faster lifestyle in the water flea Daphnia. Funct. Ecol. 32: 2225–2240.

Brans, K. I., M. Jansen, J. Vanoverbeke, N. Tiizun, R. Stoks, and L. De Meester. 2017. The heat is on: Genetic adaptation to urbanization mediated by thermal tolerance and body size. Glob. Chang. Biol. 23: 5218–5227.

Caizergues, A. E., J. S. Santangelo, R. W. Ness, F. Angeoletto, D. N. Anstett, J. Anstett, F. Baena-Diaz, E. J. Carlen, J. A. Chaves, M. S. Comerford, K. Dyson, M. Falahati-Anbaran, M. D. E. Fellowes, K. A. Hodgins, G. R. Hood, C. Ifiiguez­Armijos, N. J. Kooyers, A. Lazaro-Lobo, A. T. Moles, J. Munshi-South, J. Paule, I. M. Porth, L. Y. Santiago-Rosario, K. S. Whitney, A. J. M. Tack, and M. T. J. Johnson. 2024. Does urbanisation lead to parallel demographic shifts across the world in a cosmopolitan plant? Mal. Ecol. 33: el 7311.

Campbell-Staton, S. C., K. M. Winchell, N. C. Rochette, J. Fredette, I. Maayan, R. M. Schweizer, and J. Catchen. 2020. Parallel selection on thermal physiology facilitates repeated adaptation of city lizards to urban heat islands. Nat Ecol Evol 4: 652–658.

Canada. Dept. of Agriculture. 1941. White clover in Canada. Tech. rep. Ottawa: Experimental Farms Service. Forage Crops Division.

Carlen, E. and J. Munshi-South. 2021. Widespread genetic connectivity of feral pigeons across the Northeastern megacity. Evol. Appl. 14: 150–162.

Chen, J., J. Yu, L. Ge, H. Wang, A. Berbel, Y. Liu, Y. Chen, G. Li, M. Tadege, J. Wen, V. Cosson, K. S. Mysore, P. Ratet, F. Madueno, G. Bai, and R. Chen. 2010. Control of dissected leaf morphology by a Cys(2)His(2) zinc finger transcription factor in the model legume *Medicago truncatula*. Proc. Natl. Acad. Sci. U. S. A. 107: 10754–10759.

Chen, S., Y. Zhou, Y. Chen, and J. Gu. 2018. fastp: an ultra-fast all-in-one FASTQ preprocessor. Bioinformatics 34: i884–i890.

Combs, M., K. Byers, B. M. Ghersi, M. J. Blum, A. Caccone, F. Costa, C. G. Himsworth, J. L. Richardson, and J. Munshi-South. 2018. Urban rat races: spa­ tial population genomics of brown rats (Rattus norvegicus) compared across mul­ tiple cities. Proceedings of the Royal Society B: Biological Sciences 285: 20180245.

Cvetkovic, J., I. Haferkamp, R. Rode, I. Keller, B. Pommerrenig, O. Trentmann, J. Altensell, M. Fischer-Stettler, S. Eicke, S. C. Zeeman, and H. E. Neuhaus. 2021. Ectopic maltase alleviates dwarf phenotype and improves plant frost tolerance of maltose transporter mutants. Plant Physiol. 186: 315–329.

Daday, H. 1954. Gene frequencies in wild populations of *Trifolium repens* I. Distri­ bution by latitude. Heredity 8: 61–78.

Daday, H. 1965. Gene frequencies in wild populations of *Trifolium repens* L IV. Mechanism of natural selection. Heredity 20: 355–365.

Danecek, P., J. K. Bonfield, J. Liddle, J. Marshall, V. Ohan, M. O. Pollard, A. Whitwham, T. Keane, S. A. McCarthy, R. M. Davies, and H. Li. 2021. Twelve years of SAMtools and BCFtools. Gigascience 10: giab008.

De Lucas, J. A., J. W. Forster, K. F. Smith, and G. C. Spangenberg. 2012. Assess­ ment of gene flow in white clover (*Trifolium repens* L.) under field conditions in Australia using phenotypic and genetic markers. Crop Pasture Sci. 63: 155–163.

Delaneau, O., J.-F. Zagury, M. R. Robinson, J. L. Marchini, and E.T. Dermitzakis. 2019. Accurate, scalable and integrative haplotype estimation. Nat. Commun. 10: 5436.

Des Roches, S., K. I. Brans, M. R. Lambert, L. R. Rivkin, A. M. Savage, C. J. Schell, C. Correa, L. De Meester, S. E. Diamond, N. B. Grimm, N. C. Harris, L. Govaert, A. P. Hendry, M. T. J. Johnson, J. Munshi-South, E. P. Palkovacs, M. Szulkin, M. C. Urban, B. C. Verrelli, and M. Alberti. 2020. Socio-eco-evolutionary dynamics in cities. Evol. Appl. 14: 248–267.

Diamond, S. E., L. Chick, A. B. E. Perez, S. A. Strickler, and R. A. Martin. 2017. Rapid evolution of ant thermal tolerance across an urban-rural temperature cline. Biol. J. Linn. Soc. Land. 121: 248–257.

Diamond, S. E., L. D. Chick, A. Perez, S. A. Strickler, and R. A. Martin. 2018. Evolution of thermal tolerance and its fitness consequences: parallel and non­ parallel responses to urban heat islands across three cities. Proceedings of the Royal Society B: Biological Sciences 285: 20180036.

Diamond, S. E. and R. A. Martin. 2021. Evolution in Cities. Annu. Rev. Ecol. Evol. Syst.

Ellis, E. C. and N. Ramankutty. 2008. Putting people in the map: anthropogenic biomes of the world. Front. Ecol. Environ. 6: 439–447.

Evanno, G., S. Regnaut, and J. Gaudet. 2005. Detecting the number of clusters of individuals using the software STRUCTURE: a simulation study. Mal. Ecol. 14: 2611–2620.

Evans, K. L., J. Newton, K. J. Gaston, S. P. Sharp, A. Mcgowan, and B. J. Hatchwell. 2012. Colonisation of urban environments is associated with reduced migratory behaviour, facilitating divergence from ancestral populations. Oikos 121: 634–640.

Ewels, P., M. Magnusson, S. Lundin, and M. Kaller. 2016. MultiQC: summarize analysis results for multiple tools and samples in a single report. Bioinformatics 32: 3047–3048.

Ferrer-Admetlla, A., M. Liang, T. Korneliussen, and R. Nielsen. 2014. On detecting incomplete soft or hard selective sweeps using haplotype structure. Mal. Biol. Evol. 31: 1275–1291.

Foley, J. A., R. Defries, G. P. Asner, C. Barford, G. Bonan, S. R. Carpenter, F. S. Chapin, M. T. Coe, G. C. Daily, H.K. Gibbs, J. H. Helkowski, T. Holloway, E. A. Howard, C. J. Kucharik, C. Monfreda, J. A. Patz, I. C. Prentice, N. Ramankutty, and P. K. Snyder. 2005. Global consequences of land use. Science 309: 570–574.

Fox, E. A., A. E. Wright, M. Fumagalli, and F. G. Vieira. 2019. ngsLD: evaluating linkage disequilibrium using genotype likelihoods. Bioinformatics 35: 3855–3856.

Fusco, N. A., B. J. Cosentino, J.P. Gibbs, M. L. Allen, A. J. Blumenfeld, G. H. Boettner, E. J. Carlen, M. Collins, C. Dennison, D. DiGiacopo, A.-P. Drapeau Picard, J. Edmonson, M. C. Fisher-Reid, R. Fyffe, T. Gallo, A. Grant, W. Harbold, S. B. Heard, D. J. R. Lafferty, R. M. Lehtinen, S. Marino, J. E. McDonald, A. Mortelliti, M. Murray, A. Newman, K. N. Oswald, C. Ott-Conn, J. L. Richardson, R. Rimbach, P. Salaman, M. Steele, M. R. Stothart, M. C. Urban, K. Vandegrift, J. P. Vanek, S. N. Vanderluit, L. Vezina, and A. Caccone. 2024. Population genomic structure of a widespread, urban-dwelling mammal: The eastern grey squirrel (*Sciurus carolinensis)*. Mol. Ecol. 33: el 7230.

Garrison, E. and G. Marth. 2012. Haplotype-based variant detection from short-read sequencing. arXiv *{q-bio.GN}:* 1–9.

Garud, N. R., P. W. Messer, E. O. Buzbas, and D. A. Petrov. 2015. Recent selective sweeps in North American *Drosophila melanogaster* show signatures of soft sweeps. PLoS Genet. 11: e1005004.

Gharabli, H., V. Della Gala, and D. H. Welner. 2023. The function of UDP-glycosyltransferases in plants and their possible use in crop protection. Biotechnol. Adv. 67: 108182.

Griffiths, A. G., R. Moraga, M. Tausen, V. Gupta, P. Timothy, and A. Griffiths. 2019. Breaking free: the genomics of allopolyploidy-facilitated niche expansion in white clover. Plant Cell 31: 1466–1487.

Guo, P., L. Chong, Z. Jiao, R. Xu, Q. Niu, and Y. Zhu. 2025. Salt stress activates the CDK8-AHL10-SUVH2/9 module to dynamically regulate salt tolerance in *Arabidopsis*. Nat. Commun. 16: 2454.

Han, E., J. S. Sinsheimer, and J. Novembre. 2013. Characterizing bias in population genetic inferences from low-coverage sequencing data. Mal. Biol. Evol. 31: 723–735.

Hanghöj, K., I. Moltke, P. A. Andersen, A. Manica, and T. S. Korneliussen. 2019. Fast and accurate relatedness estimation from high-throughput sequencing data in the presence of inbreeding. Gigascience 8.

Harpak, A., N. Garud, N. A. Rosenberg, D. A. Petrov, M. Combs, P. S. Pennings, and J. Munshi-South. 2021. Genetic adaptation in New York City rats. Genome Biol. Evol. 13.

Harris, S. E. and J. Munshi-South. 2017. Signatures of positive selection and local adaptation to urbanization in white-footed mice (Peromyscus leucopus). Mal. Ecol. 26: 6336–6350.

Harris, S. E., A. T. Xue, D. Alvarado-Serrano, J. T. Boehm, T. Joseph, M. J. Hickerson, and J. Munshi-South. 2016. Urbanization shapes the demographic history of a native rodent (the white-footed mouse, *Peromyscus leucopus)* in New York City. Biol. Lett. 12: 20150983-.

Hayward, L. K. and G. Sella. 2022. Polygenic adaptation after a sudden change in environment. Elife 11.

He, M., M. Lan, B. Zhang, Y. Zhou, Y. Wang, L. Zhu, M. Yuan, and Y. Fu. 2018. Rab-Hlb is essential for trafficking of cellulose synthase and for hypocotyl growth in *Arabidopsis thaliana*. J. Integr. Plant Biol. 60: 1051–1069.

Hedrick, P. W. and R. C. Lacy. 2015. Measuring relatedness between inbred indi­ viduals. J. Hered. 106: 20–25.

Hoban, S., J. L. Kelley, K. E. Lotterhos, M. F. Antolin, G. Bradburd, D. B. Lowry, M. L. Poss, L. K. Reed, A. Storfer, and M. C. Whitlock. 2016. Finding the ge­ nomic basis oflocal adaptation: pitfalls, practical solutions, and future directions. Am. Nat. 188: 379–397.

Hodgins, K. A. and S. Yeaman. 2019. Mating system impacts the genetic architecture of adaptation to heterogeneous environments. New Phytol. 224: 1201–1214.

Hu, J., Y. Zhang, J. Wang, and Y. Zhou. 2014. Glycerol affects root development through regulation of multiple pathways in *Arabidopsis*. PLoS One 9: e86269.

Hudson, R. R., M. Slatkin, and W. P. Maddison. 1992. Estimation of levels of gene flow from DNA sequence data. Genetics 132: 583–589.

Innes, S. G., J. S. Santangelo, N. J. Kooyers, K. M. Olsen, and M. T. J. Johnson. 2022. Evolution in response to climate in the native and introduced ranges of a globally distributed plant. Evolution 76: 1495–1511.

Inostraza, L., M. Bhakta, H. Acuna, C. Vasquez, J. Ibanez, G. Tapia, W. Mei, M. Kirst, M. Resende, and P. Munoz. 2018. Understanding the complexity of cold tolerance in white clover using temperature gradient locations and a GWAS approach. Plant Genome 11: 170096.

Jia, Q., D. Kong, Q. Li, S. Sun, J. Song, Y. Zhu, K. Liang, Q. Ke, W. Lin, and J. Huang. 2019. The function of inositol phosphatases in plant tolerance to abiotic stress. Int. J. Mal. Sci. 20: 3999.

Jiang, L., Y. Fu, X. Tian, Y. Ma, F. Chen, and G. Wang. 2022. The *Anthurium* APRR2-like gene promotes photosynthetic pigment accumulation in response to salt stress. Trap. Plant Biol. 15: 12–21.

Johnson, M. T. J. and J. Munshi-South. 2017. Evolution of life in urban environ­ ments. Science 358: eaam8327.

Johnson, M. T. J., C. Prashad, M. Lavoignat, and H. S. Saini. 2018. Contrasting the effects of natural selection, genetic drift and gene flow on urban evolution in white clover (*Trifolium repens*). Proceedings of the Royal Society B: Biological Sciences 285: 20181019.

Johri, P., C. F. Aquadro, M. Beaumont, B. Charlesworth, L. Excoffier, A. Eyre­Walker, P. D. Keightley, M. Lynch, G. McVean, B. A. Payseur, S. P. Pfeifer, W. Stephan, and J. D. Jensen. 2022. Recommendations for improving statistical inference in population genomics. PLoS Biol. 20: e3001669.

Kamdem, C., C. Fouet, S. Gamez, and B. J. White. 2017. Pollutants and insecticides drive local adaptation in african malaria mosquitoes. Mal. Biol. Evol. 34: 1261–1275.

KjIBrgaard, T. 2003. A plant that changed the world: the rise and fall of clover 1000-2000. Landscape Res. 28: 41–49.

Kooyers, N. J. and K. M. Olsen. 2013. Searching for the bull’s eye: agents and targets of selection vary among geographically disparate cyanogenesis dines in white clover *(Trifolium repens* L.) Heredity 111: 495–504.

Kopelman, N. M., J. Mayzel, M. Jakobsson, N. A. Rosenberg, and I. Mayrose. 2015. Clumpak: a program for identifying clustering modes and packaging population structure inferences across K. Mal. Ecol. Resour. 15: 1179–1191.

Korneliussen, T. S., A. Albrechtsen, and R. Nielsen. 2014. ANGSD: analysis of next generation sequencing data. BMC Bioinformatics 15: 1–13.

Korunes, K. L. and K. Samuk. 2021. pixy: Unbiased estimation of nucleotide di­ versity and divergence in the presence of missing data. Mal. Ecol. Resour. 21: 1359–1368.

Kreiner, J. M., S. M. Latorre, H. A. Burbano, J. R. Stinchcombe, S. P. Otto, D. Weigel, and S. I. Wright. 2022a. Rapid weed adaptation and range expansion in response to agriculture over the past two centuries. Science 378: 1079–1085.

Kreiner, J. M., G. Sandler, A. J. Stern, P. J. Tranel, D. Weigel, J. Stinchcombe, and S. I. Wright. 2022b. Repeated origins, widespread gene flow, and allelic interactions of target-site herbicide resistance mutations. Elife 11.

Kuo, W.-H., S. J. Wright, L. L. Small, and K. M. Olsen. 2024a. De novo genome assembly of white clover (*Trifolium repens* L.) reveals the role of copy number variation in rapid environmental adaptation. BMC Biol. 22: 165.

Kuo, W.-H., L. Zhong, S. J. Wright, D. M. Goad, and K. M. Olsen. 2024b. Beyond cyanogenesis: Temperature gradients drive environmental adaptation in North American white clover (Trifolium repens L.) Mal. Ecol.: el 7484.

Lasky, J. R., E. B. Josephs, and G. P. Morris. 2023. Genotype-environment associ­ ations to reveal the molecular basis of environmental adaptation. Plant Cell 35: 125–138.

Lau, J. A. and J. L. Funk. 2023. How ecological and evolutionary theory expanded the ‘ideal weed’ concept. Oecologia 203: 251–266.

Lenormand, T. 2002. Gene flow and the limits to natural selection. Trends Ecol. Evol. 17: 183–189.

Li, H. 2011a. A statistical framework for SNP calling, mutation discovery, association mapping and population genetical parameter estimation from sequencing data. Bioinformatics 27: 2987–2993.

Li, H. 2011b. Improving SNP discovery by base alignment quality. Bioinformatics 27: 1157–1158.

Li, H. and R. Durbin. 2009. Fast and accurate short read alignment with Burrows­ Wheeler transform. Bioinformatics 25: 1754–1760.

Li, H., B. Handsaker, A. Wysoker, T. Fennell, J. Ruan, N. Homer, G. Marth, G. Abecasis, R. Durbin, and 1000 Genome Project Data Processing Subgroup. 2009. The Sequence Alignment/Map format and SAMtools. Bioinformatics 25: 2078–2079.

Littleford-Colquhoun, B. L., C. Clemente, M. J. Whiting, D. Ortiz-Barrientos, and C. H. Frere. 2017. Archipelagos of the Anthropocene: rapid and extensive differ­ entiation of native terrestrial vertebrates in a single metropolis. Mal. Ecol. 26: 2466–2481.

Løjtnant, C. L., B. Boelt, S. K. Clausen, C. Damgaard, P. Kryger, A. Novy, M. Philipp, C. H. Ingvordsen, and R. B. Jørgensen. 2012. Modelling gene flow be­ tween fields of white clover with honeybees as pollen vectors. Environ. Model. Assess. 17: 421–430.

Lotterhos, K. E. and M. C. Whitlock. 2015. The relative power of genome scans to detect local adaptation depends on sampling design and statistical method. Mal. Ecol. 24: 1031–1046.

Lourerno, A., D. Alvarez, I. J. Wang, and G. Velo-Anton. 2017. Trapped within the city: integrating demography, time since isolation and population-specific traits to assess the genetic effects of urbanization. Mol. Ecol. 26: 1498–1514.

Martin, E., S. El-Galmady, and M. T. J. Johnson. 2024. Urban socioeconomic vari­ ation influences the ecology and evolution of trophic interactions. Ecol. Lett. 27: el 4407.

Martin, M., M. Patterson, S. Garg, S. O. Fischer, N. Pisanti, G. W. Klau, A. Schoenhuth, and T. Marschall. 2016. WhatsHap: fast and accurate read-based phasing. bioRxiv: 085050.

Martin, R. A., L. D. Chick, M. L. Garvin, and S. E. Diamond. 2021. In a nutshell, a reciprocal transplant experiment reveals local adaptation and fitness trade-offs in response to urban evolution in an acorn-dwelling ant. Evolution 75: 876–887.

Meisner, J. and A. Albrechtsen. 2018. Inferring population structure and admixture proportions in low-depth NGS data. Genetics 210: 719–731.

Miles, L. S., L. R. Rivkin, M. T. J. Johnson, J. Munshi-South, and B. C. Verrelli. 2019. Gene flow and genetic drift in urban environments. Mal. Ecol. 28: 4138–4151.

Molder, F., K. P. Jablonski, B. Letcher, M. B. Hall, C.H. Tomkins-Tinch, V. Sochat, J. Forster, S. Lee, S. O Twardziok, A. Kanitz, A. Wilm, M. Holtgrewe, S. Rahmann, S. Nahnsen, and J. Koster. 2021. Sustainable data analysis with Snake­ make. Fl 000Res. 10: 33.

Molesini, B., S. Zanzoni, G. Mennella, G. Francese, A. Losa, G. L Rotino, and T. Pandolfini. 2017. The *Arabidopsis* N-acetylornithine deacetylase controls or­ nithine biosynthesis via a linear pathway with downstream effects on polyamine levels. Plant Cell Physiol. 58: 130–144.

Mueller, J., K. Heiner, S. Boerno, J. Tella, M. Carrete, and B. Kempenaers. 2018. Evolution of genomic variation in the burrowing owl in response to recent colo­ nization of urban areas. Proceedings of the Royal Society B: Biological Sciences 285: 20180206.

Mueller, J. C., M. Carrete, S. Boerno, H. Kuhl, J. L. Tella, and B. Kempenaers. 2020. Genes acting in synapses and neuron projections are early targets of selection during urban colonization. Mal. Ecol. 29: 3403–3412.

Munshi-South, J. 2012. Urban landscape genetics: Canopy cover predicts gene flow between white-footed mouse *(Peromyscus leucopus)* populations in New York City. Mal. Ecol. 21: 1360–1378.

Munshi-South, J. and K. Kharchenko. 2010. Rapid, pervasive genetic differentiation of urban white-footed mouse *(Peromyscus leucopus)* populations in New York City. Mal. Ecol. 19Munshi-S: 4242–4254.

Neve, P., M. Vila-Aiub, and F. Roux. 2009. Evolutionary-thinking in agricultural weed management. New Phytol. 184: 783–793.

Newbold, T., L. N. Hudson, S. L. L. Hill, S. Contu, I. Lysenko, R. A. Senior, L. Borger, D. J. Bennett, A. Choimes, B. Collen, J. Day, A. De Palma, S. Diaz, S. Echeverria-Londono, M. J. Edgar, A. Feldman, M. Garon, M. L. K. Harrison, T. Alhusseini, D. J. Ingram, Y. Itescu, J. Kattge, V. Kemp, L. Kirkpatrick, M. Kleyer, D. L. P. Correia, C. D. Martin, S. Meiri, M. Novosolov, Y. Pan, H. R. P. Phillips, D. W. Purves, A. Robinson, J. Simpson, S. L. Tuck, E. Weiher, H. J. White, R. M. Ewers, G. M. Mace, J. P. W. Scharlemann, and A. Purvis. 2015. Global effects of land use on local terrestrial biodiversity. Nature 520: 45–50.

Nielsen, R., T. Korneliussen, A. Albrechtsen, Y. Li, and J. Wang. 2012. SNP calling, genotype calling, and sample allele frequency estimation from new-generation sequencing data. PLoS One 7: e37558.

Nielsen, R., J. S. Paul, A. Albrechtsen, and Y. S. Song. 2011. Genotype and SNP calling from next-generation sequencing data. Nat. Rev. Genet. 12: 443–451.

Noel, S., M. Ouellet, P. Galois, and F. J. Lapointe. 2007. Impact of urban fragmen­ tation on the genetic structure of the eastern red-backed salamander. Conserv. Genet. 8: 599–606.

Okonechnikov, K., A. Conesa, and F. Garcia-Alcalde. 2016. Qualimap 2: advanced multi-sample quality control for high-throughput sequencing data. Bioinformat­ ics 32: 292–294.

Olsen, K. M., D. M. Goad, S. J. Wright, M. L. Dutta, S. R. Myers, L. L. Small, and L.-F. Li. 2021. Dual-species origin of an adaptive chemical defense polymorphism. New Phytol. 232: 1477–1487.

Oziolor, E. M., N. M. Reid, S. Yair, K. M. Lee, S. G. Verploeg, P. C. Bruns, J. R. Shaw, A. Whitehead, and C. W. Matson. 2019. Adaptive introgression enables evolutionary rescue from extreme environmental pollution. Science 126: 455–457.

Pickering, C. and A. Mount. 2010. Do tourists disperse weed seed? A global review of unintentional human-mediated terrestrial seed dispersal on clothing, vehicles and horses. J. Sustainable Tour. 18: 239–256.

Prileson, E. G. and R. A. Martin. 2024. Evolution and plasticity of physiological traits in the collembolan *Orchesella villosa* at fine spatial scales within the city. Biol. J. Linn. Soc. Land.: blae038.

Pullagurla, N. J., S. Shome, R. Yadav, and D. Laha. 2023. ITPKl regulatesjasmonate­ controlled root development in *Arabidopsis Thaliana*. Biomolecules 13: 1368.

Pya, N. and S. N. Wood. 2015. Shape constrained additive models. Stat. Comput. 25: 543–559.

R Core Team. 2020. R: A language and environment for statistical computing. Version 3.6.3. Vienna, Austria.

Ravinet, M., T. Elgvin, C. Trier, M. Aliabadian, A. Gavrilov, and G.-P. Saetre. 2018. Signatures of human-commensalism in the house sparrow genome. Proceedings of the Royal Society B: Biological Sciences 285: 20181246.

Reid, N. M., D. A. Proestou, B. W. Clark, W. C. Warren, J. K. Colbourne, J. R. Shaw, S. I. Karchner, M. E. Hahn, D. Nacci, M. F. Oleksiak, D. L. Crawford, and A. Whitehead. 2016. The genomic landscape of rapid repeated evolutionary adaptation to toxic pollution in wild fish. Science 354: 1305 LP–1308.

Rivkin, L. R. and M. T. J. Johnson. 2022. The impact of urbanization on outcrossing rate and population genetic variation in the native wildflower, *Impatiens capensis*. J Urban Ecol 8.

Salmon, P., A. Jacobs, D. Ahren, C. Biard, N. J. Dingemanse, D. M. Dominoni, B. Helm, M. Lundberg, J. C. Senar, P. Sprau, M. E. Visser, and C. Isaksson. 2021. Continent-wide genomic signatures of adaptation to urbanisation in a songbird across Europe. Nat. Commun. 12: 2983.

Santangelo, J. S., P. Battlay, B. T. Hendrickson, W.-H. Kuo, K. M. Olsen, N. J. Kooyers, M. T. J. Johnson, K. A. Hodgins, and R. W. Ness. 2023. Haplotype­ Resolved, Chromosome-Level Assembly of White Clover (*Trifolium repens* L., Fabaceae). Genome Biol. Evol. 15.

Santangelo, J. S., L. R. Rivkin, C. Advenard, and K. A. Thompson. 2020a. Mul­ tivariate phenotypic divergence along an urbanization gradient. Biol. Lett. 16: 20200511.

Santangelo, J. S., L. R. Rivkin, and M. T. J. Johnson. 2018. The evolution of city life. Proceedings of the Royal Society B: Biological Sciences 285: 20181529.

Santangelo, J. S., C. Roux, and M. T. J. Johnson. 2022a. The effects of environmental heterogeneity within a city on the evolution of dines. J. Ecol.

Santangelo, J. S., K. A. Thompson, B. Cohan, J. Syed, R. W. Ness, and M. T. J. Johnson. 2020b. Predicting the strength of urban-rural dines in a Mendelian polymorphism. Evolution Letters 4: 212–225.

Santangelo, J. S. et al. 2022b. Global urban environmental change drives adaptation in white clover. Science 375: 1275–1281.

Schmidt, C., M. Domaratzki, R. P. Kinnunen, J. Bowman, and C. J. Garroway. 2020. Continent-wide effects of urbanization on bird and mammal genetic diversity. Proc. Biol. Sci. 287: 20192497.

Shockey, J. and J. Browse. 2011. Genome-level and biochemical diversity of the acyl­ activating enzyme superfamily in plants: Biochemistry and evolution of plant AAE proteins. Plant J. 66: 143–160.

Sincik, M. and E. Acikgoz. 2007. Effects of white clover inclusion on turf character­ istics, nitrogen fixation, and nitrogen transfer from white clover to grass species in turf mixtures. Commun. Soil Sci. Plant Anal. 38: 1861–1877.

Skotte, L., T. S. Korneliussen, and A. Albrechtsen. 2013. Estimating individual admixture proportions from next generation sequencing data. Genetics 195: 693–702.

Snaydon, R. W. and M. S. Davies. 1972. Rapid population differentiation in a mosiac environment II: Morphological variation in *Anthoxanthum odoratum*. Evolution 26: 390–405.

Szpiech, Z. A. and R. D. Hernandez. 2014. selscan: an efficient multithreaded pro­ gram to perform EHR-based scans for positive selection. Mal. Biol. Evol. 31: 2824–2827.

Szpiech, Z. A., T. E. Novak, N. P. Bailey, and L. S. Stevison. 2021. Application of a novel haplotype-based scan for local adaptation to study high-altitude adaptation in rhesus macaques. Evol. Lett. 5: 408–421.

Szulkin, M., J. Munshi-South, and A. Charmantier. 2020. Urban Evolutionary Biology. Oxford University Press, Oxford, United Kingdom.

Taylor, N. L. 2008. A century of clover breeding developments in the United States. Crop Sci. 48: 1–13.

Theodorou, P., R. Radzeviciute, B. Kahnt, A. Soro, I. Grosse, and R. J. Paxton. 2018. Genome-wide single nucleotide polymorphism scan suggests adaptation to urbanization in an important pollinator, the red-tailed bumblebee *(Bombus lapi­ darius* L.) Proceedings of the Royal Society B: Biological Sciences 285: 20172806.

Thompson, K. A., M. Renaudin, and M. T. J. Johnson. 2016. Urbanization drives the evolution of parallel dines in plant populations. Proceedings of the Royal Society B: Biological Sciences 283: 20162180.

Torres, R., Z. A. Szpiech, and R. D. Hernandez. 2018. Human demographic history has amplified the effects of background selection across the genome. PLoS Genet. 14: e1007387.

Tsitsekian, D., G. Daras, A. Alatzas, D. Templalexis, P. Hatzopoulos, and S. Rigas. 2019. Comprehensive analysis of Lon proteases in plants highlights independent gene duplication events. J. Exp. Bot. 70: 2185–2197.

Tsujimoto, Y., T. Numaga, K. Ohshima, M.-A. Yano, R. Ohsawa, D. B. Goto, S. Naito, and M. Ishikawa. 2003. *Arabidopsis* TOBAMOVIRUS MULTIPLICA­ TION (TOM) 2 locus encodes a transmembrane protein that interacts with TOMI. EMBO J. 22: 335–343.

Turcotte, M. M., H. Araki, D. S. Karp, K. Poveda, and S. R. Whitehead. 2017. The eco-evolutionary impacts of domestication and agricultural practices on wild species. Philos. Trans. R. Soc. Land. B Biol. Sci. 372.

Unfried, T. M., L. Hauser, and J. M. Marzluff. 2013. Effects of urbanization on Song Sparrow *(Melospiza melodia)* population connectivity. Conserv. Genet. 14: 41–53.

Vasimuddin, M., S. Misra, H. Li, and S. Aluru. 2019. Efficient architecture-aware acceleration of BWA-MEM for multicore systems. In: 2019 IEEE International Parallel and Distributed Processing Symposium (IPDPS): pp. 314–324.

Verrelli, B. C., M. Alberti, S. Des Roches, N. C. Harris, A. P. Hendry, M. T. J. Johnson, A. M. Savage, A. Charmantier, K. M. Gotanda, L. Govaert, L. S. Miles, L. R. Rivkin, K. M. Winchell, K. I. Brans, C. Correa, S. E. Diamond, B. Fitzhugh, N. B. Grimm, S. Hughes, J. M. Marzluff, J. Munshi-South, C. Rojas, J. S. Santangelo, C. J. Schell, J. A. Schweitzer, M. Szulkin, M. C. Urban, Y. Zhou, and C. Ziter. 2022. A global horizon scan for urban evolutionary ecology. Trends Ecol. Evol.

Voight, B. F., S. Kudaravalli, X. Wen, and J. K. Pritchard. 2006. A map of recent positive selection in the human genome. PLoS Biol. 4: e72.

Wang, X., J. Ding, S. Lin, D. Liu, T. Gu, H. Wu, R. N. Trigiano, R. McAvoy, J. Huang, and Y. Li. 2020. Evolution and roles of cytokinin genes in angiosperms 2: Do ancient CKXs play housekeeping roles while non-ancient CKXs play reg­ ulatory roles? Hortic. Res. 7: 29.

Weigand, H. and F. Leese. 2018. Detecting signatures of positive selection in non­ model species using genomic data. Zool J Linn Soc 184: 528–583.

Whiting, J. R. 2022. Jim Whiting91/genotype_plot: Genotype Plot. Version v0.2.1.

Winchell, K. M., I. Maayan, J. Fredette, and L. J. Revell. 2018. Linking locomotor performance to morphological shifts in urban lizards. Proceedings of the Royal Society Biological Sciences 285: 20180229.

Winchell, K. M., E. J. Carlen, A. R. Puente-Rolon, and L. J. Revell. 2017. Divergent habitat use of two urban lizard species. Ecol. Evol. 8: 25–35.

Winchell, K. M., R. G. Reynolds, S. R. Prado-Irwin, A. R. Puente-Rolon, and L. J. Revell. 2016. Phenotypic shifts in urban areas in the tropical lizard *Anolis cristatellus*. Evolution 70: 1009–1022.

Wu, R., W. Zheng, J. Tan, R. Sammer, L. Du, and C. Lu. 2019. Protein partners of plant ubiquitin-specific proteases (UBPs). Plant Physiol. Biochem. 145: 227–236.

Yakub, M. and P. Tiffin. 2017. Living in the city: urban environments shape the evolution of a native annual plant. Glob. Chang. Biol. 23: 2082–2089.

Yamaguchi, N., S. Matsubara, K. Yoshimizu, M. Seki, K. Hamada, M. Kamitani, Y. Kurita, Y. Nomura, K. Nagashima, S. Inagaki, T. Suzuki, E.-S. Gan, T. To, T. Kakutani, A. J. Nagano, A. Satake, and T. Ito. 2021. H3K27me3 demethy­ lases alter HSP22 and HSPl 7.6C expression in response to recurring heat m *Arabidopsis*. Nat. Commun. 12: 3480.

Yeaman, S. 2022. Evolution of polygenic traits under global vs local adaptation. Genetics 220.

